# A zebrafish ribosomopathy links Ddx49-dependent ribosome biogenesis to differential failure of population expansion among pigment cell lineages

**DOI:** 10.64898/2026.10.13.758983

**Authors:** Yipeng Liang, Kinga Malezyna, Abigail J Dennis, Abhishek Kumar, Braedan M McCluskey, Diane S Sepich, Liliana Solnica-Krezel, Arunkumar Dhayalan, Ahmad Jomaa, Larissa B Patterson, David M Parichy

## Abstract

Defects in ribosome biogenesis cause surprisingly cell-type specific pathologies. Why some cells are more vulnerable than others remains unresolved. Here, we show that a zebrafish pigment pattern mutant, *sisley*, results from a viable missense allele of *ddx49*, encoding a DEAD-box RNA helicase required for small ribosomal subunit biogenesis and early survival. In mutants, yellow/orange xanthophores fail to proliferate during development of the adult stripe pattern, whereas closely related melanophores are affected only mildly. The mutant protein has reduced helicase but increased ATPase activity and, consistent with impaired ribosome biogenesis *in vivo*, mutants are deficient in 18S rRNA relative to its precursor. Remarkably, a similar deficiency in 18S rRNA relative to precursor rRNA distinguishes xanthophores from melanophores even in the wild-type, suggesting an inherent difference in rRNA processing that leaves xanthophores vulnerable to Ddx49 dysfunction. Both xanthophore deficiency and a male-biased sex ratio of the *sisley* mutant were rescued by mutation of *tp53*. Our findings suggest how baseline differences in ribosome production shape cell-type specific outcomes of mutations affecting ribosome biogenesis, and provide an adult-viable model for Ddx49 function *in vivo*.

## Introduction

Pigmentation of vertebrates has long been a useful system for interrogating fundamental mechanisms of pattern formation and how differentiation and morphogenesis at the level of individual cells translate into phenotypes that are often species-specific (Weller et al., 2026). Pigment patterns of mammals and birds arise from widely distributed neural crest derived melanoblasts that differentiate in spatially discrete domains as melanocytes, which then deliver melanin to keratinocytes for incorporation into developing hair or feathers. By contrast, the pigment patterns of amphibians, reptiles and fishes depend on the numbers and arrangements of several classes of neural crest derived chromatophores that retain pigments intracellularly, including melanophores (homologues of melanocytes), xanthophores with yellow or orange pigments, and iridophores with reflective platelets of crystalline guanine (Parichy, 1996; Irion et al., 2016; Manukyan et al., 2017; Parichy, 2021; Helleboid and Tzika, 2026). In zebrafish, for example, dark stripes of melanophores and sparse iridophores alternate with light interstripes of xanthophores and densely packed iridophores, in a pattern that forms during the larva-to-adult transition (Patterson and Parichy, 2019). Patterns often depend on these cells organizing as they are differentiating, choreographed in part by the tissue environment and in part by interactions between chromatophores themselves. Critical, too, is expansion of chromatophore populations, coordinated with one another and with often substantial somatic growth: decreases or increases in any one chromatophore class can have cascading effects, leading to very different patterns (Parichy et al., 2000; Frohnhofer et al., 2013; Patterson et al., 2014; Gur et al., 2020). Such sensitivity to cell numbers and interactions, combined with the genetic tractability of zebrafish, makes the adult pigment pattern ideal for forward genetic screens of overtly visible phenotypes

Spontaneous and induced pigmentation variants in both endotherms and ectotherms have allowed the discovery of genes essential to cell–cell communication during pattern formation (Geissler et al., 1988; Hosoda et al., 1994; Watanabe et al., 2006; Irion et al., 2014; Eom et al., 2021). They have also revealed genes and classes of genes that contribute to fundamental cellular processes across diverse tissue types and organ systems (Mercer et al., 1991; Ollmann et al., 1997; Zhang et al., 2003; Ng et al., 2009; Krauss et al., 2013). One such process is ribosome biogenesis, affected in several mouse pigmentation mutants (Oliver et al., 2004; McGowan et al., 2008; Terzian et al., 2011; Watkins-Chow et al., 2013). Assembly of small (40S) and large (60S) ribosomal subunits, largely elucidated through studies of yeast , is an exceptionally complex process involving transcription of a 47S precursor (pre-rRNA) that is processed to furnish mature ribosomal RNAs (rRNAs) to each subunit (18S; 5.8S and 28S, respectively), numerous ribosomal proteins that function within each mature subunit (33 and 47 proteins of 40S and 60S, respectively), and a veritable army of processing and accessory factors (Bassler and Hurt, 2019; Bohnsack and Bohnsack, 2019; Dorner et al., 2023; Vanden Broeck and Klinge, 2024). Beginning in nucleoli, ribosome assembly continues in the nucleoplasm and is completed in the cytoplasm. Not surprisingly, mutations of genes encoding ribosomal proteins and accessory factors are most often early lethal when homozygous, and frequently have dominant phenotypes owing to haploinsufficiency (Amsterdam et al., 2004; Marygold et al., 2007; Terzian and Box, 2013). In humans, mutations affecting rRNA abundances, ribosomal proteins, or ribosome assembly factors cause “ribosomopathies” (Danilova and Gazda, 2015; Yelick and Trainor, 2015; Farley-Barnes et al., 2019; Kampen et al., 2020; Ni and Buszczak, 2023). These disorders often have surprising specificities for cell and tissue types, given the ubiquity of ribosomes, and many depend on Tp53-mediated responses to nucleolar stress; how such specificity arises remains incompletely understood (Danilova and Gazda, 2015; Mills and Green, 2017).

Among the factors required for rRNA processing and ribosome biogenesis are DEAD-box RNA helicases (Linder and Jankowsky, 2011; Bohnsack et al., 2023; Dorner and Hondele, 2024). These enzymes, named for one of several conserved amino acid sequence motifs, contain two helicase domains similar to bacterial recombinase A (RecA1, RecA2), joined by a flexible interdomain linker that allows opening and closing around an RNA substrate. These helicases do not target specific RNA sequence motifs; rather association of the enzyme with double-stranded RNA involves interactions between the RecA-like domains and the sugar-phosphate backbone. Once bound, the enzymes exploit dynamic fluctuations in RNA structure to separate strands, initially within the helicase-bound region but propagating beyond this site without helicase translocation. Protein conformational changes that allow stable association with RNA and strand separation require the helicase to bind ATP, whereas efficient dissociation from RNA and substrate turnover require ATP hydrolysis, so that each productive cycle uses at least one ATP (Linder and Jankowsky, 2011). With their cofactors, DEAD-box helicases participate in a wide range of cellular processes, including transcription, pre-mRNA splicing, RNA export and storage, translation, nonsense-mediated RNA decay, stress responses and pre-rRNA cleavage to form mature rRNAs. Mutations in DEAD-box helicase genes are associated with pathologies of the craniofacial and limb skeleton, central nervous system, kidney, eye and other traits (Bohnsack et al., 2023). DDX49 and its yeast orthologue Dbp8 have received far less attention than many family members but have been implicated in pre-rRNA processing and small subunit assembly in cultured cells and yeast (Daugeron and Linder, 2001; Awasthi et al., 2018; Tse et al., 2026); roles in an animal have not been examined.

Here, we identify a homozygous viable missense substitution in the gene encoding DEAD-box helicase Ddx49 as the cause of an adult pigment pattern mutant phenotype in zebrafish. Mutants have a severe defect in xanthophore population expansion, with impaired formation of the adult stripe pattern. Single-cell transcriptomics reveals dysregulation of genes functioning in ribosome biogenesis and translation, with xanthophores more severely affected than other chromatophores and exhibiting a concomitant induction of nucleolar stress response genes and Tp53 target genes. The mutant protein is reduced in abundance and has discordant enzymatic activities, with helicase activity diminished but ATPase activity increased, and mutants are deficient in pre-rRNA processing; loss-of-function mutations likewise impair processing, and further lead to loss of the small ribosomal subunit and early lethality. Remarkably, even in wild-type fish, xanthophores have lower ratios of mature 18S rRNA to pre-rRNA than melanophores, suggesting that a pre-existing difference in rRNA processing contributes to their greater vulnerability. Finally, we show that both the pigment pattern defect and another phenotype, male-biased sexual differentiation, are rescued by simultaneous mutation of *tp53*, as observed for ribosomopathy phenotypes in other contexts. The zebrafish missense allele provides new insights into the cell-type specificity of ribosome dysfunction and, to our knowledge, the first *in vivo* model for interrogating *ddx49* function in an animal.

## Results

### Disrupted adult stripe formation in *stl729* mutant zebrafish

Recovered in a forward genetic screen for ethylnitrosourea-induced mutations in the AB genetic background, the *stl729* mutant allele is recessive with homozygous adults exhibiting disorganized patterns of adult stripes reminiscent of—though more variable than—xanthophore-deficient mutants for *colony stimulating factor 1 receptor a* (*csf1ra*) (Parichy et al., 2000; Parichy and Turner, 2003a) (**Figure 1A**, **Figure 1—figure supplement 1**). Melanophore stripes have gaps and poorly formed borders on the body and are variably positioned, or missing entirely, in caudal and anal fins. Adults are also deficient for yellow coloration. Embryos and early larvae (3–5 days post-fertilization) were indistinguishable from the wild-type. In keeping with prior naming conventions, we christened the mutant *sisley* (*ssl*), for impressionist landscape painter Alfred Sisley, recognized for his muted palette and rendering of varied conditions in single scenes (Shone, 1999). Reminiscent of Alfred Sisley’s failure to thrive financially, *ssl* homozygotes tend to be slow growing and developing, suggesting additional, pleiotropic defects. Homozygotes are also severely sex-biased, with *ssl/ssl* males far outnumbering females (∼500:1), as observed during routine maintenance and propagation of this allele.

**Figure 1.**
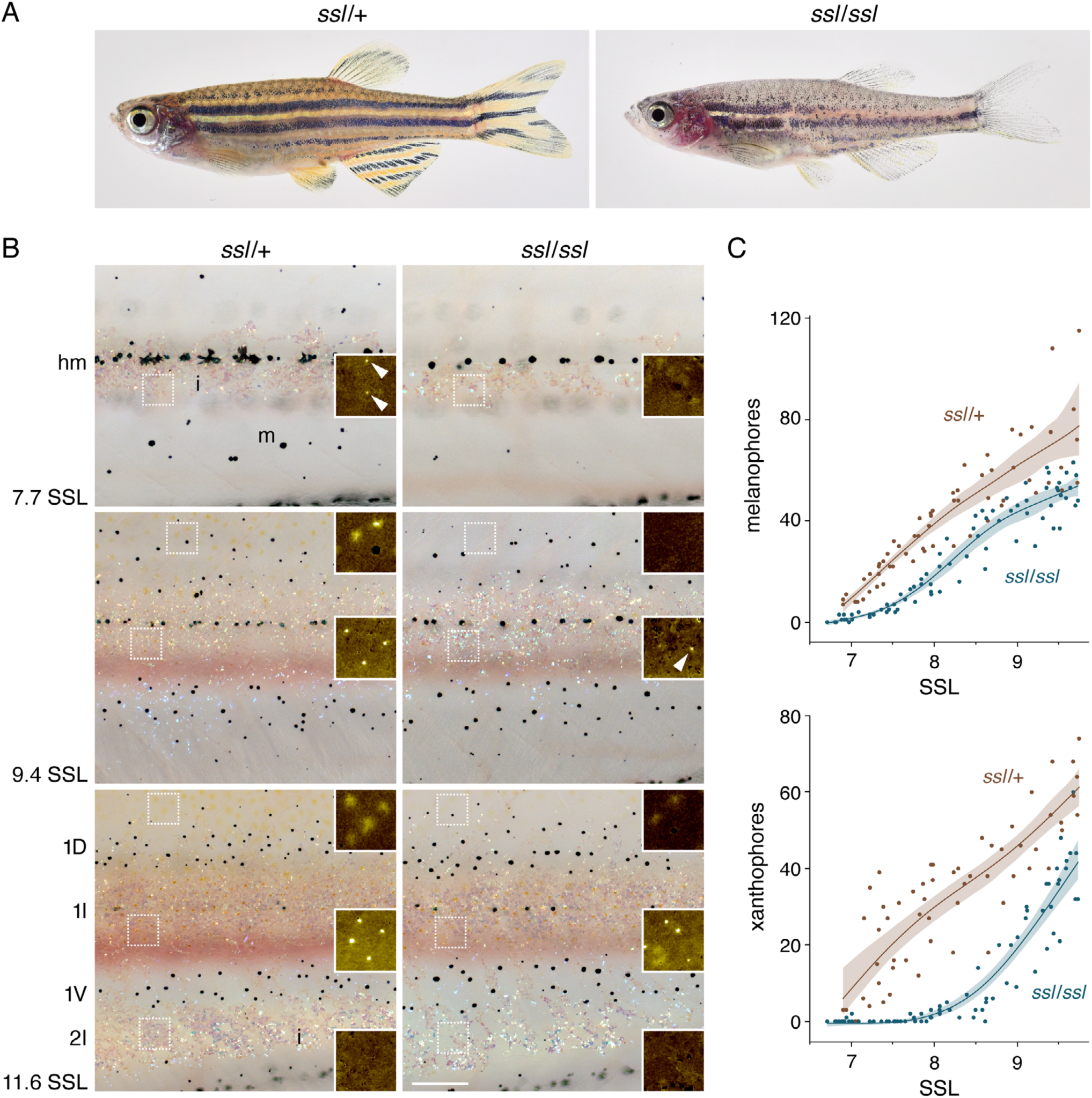
*ssl* mutants have disrupted adult stripes and defects in chromatophore complements. (**A**) Young adult fish showing well-ordered pattern of wild-type, with normal yellow coloration, and disorganized pattern of mutant, deficient for yellow coloration. (**B**) Representative images from wild-type (*ssl/+*) and mutant (*ssl/ssl*) siblings imaged daily (*n*=4 fish per genotype) through adult pattern formation and shown here when differences had already become evident (stages at lower left of each row). In these images, pigments have been contracted towards cell centers to facilitate cell counts and carotenoid pigments of xanthophores are pseudocolored in insets (e.g., arrows) for easier visualization of these cells (Korzeniwsky et al., 2025). At each stage, *ssl* mutants had fewer xanthophores and melanophores (m) than wild-type. At later stages (bottom), major pattern elements had formed but were less organized in *ssl* with defects in melanophore and xanthophore distributions as well as iridescent iridophore (i) coverage. 1D, primary dorsal stripe; 1I, primary interstripe; 1V, primary ventral stripe; 2I, secondary (ventral) interstripe; hm, position of horizontal myoseptum. Stages are indicated as standardized standard length in mm (SSL). (**C**) Numbers of melanophores (upper) and pigmented xanthophores (lower) showing especially severe deficiencies for the latter. Splines indicate trends over developmental time pooled across individuals. Scale bar, 200 µm (**B**). (Numerical data for display items here and below can be found in **Supplementary File 2 —Figure source data**.)

**Figure 1—figure supplement 1.**
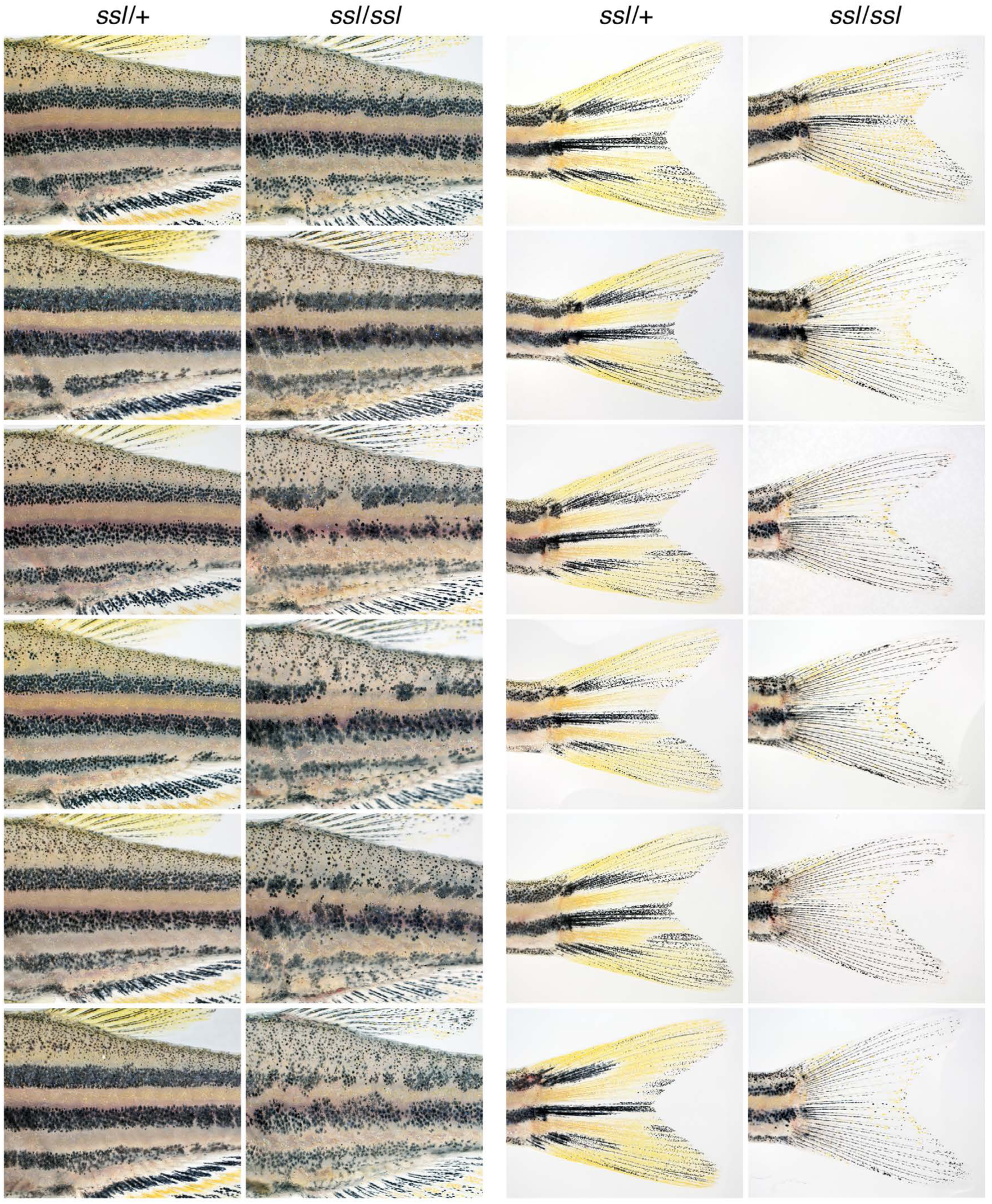
Variability of pattern defects in *ssl* mutant zebrafish. Shown are patterns on body and fins of *ssl* mutants and wild-type siblings, drawn from a single family. Fish are 15 SSL.

To understand the ontogeny of pattern defects, we imaged individually reared, phenotypically wild-type heterozygous (*ssl/+*) and homozygous mutant (*ssl/ssl*) siblings daily during adult pattern formation; hereafter, “wild-type” refers to phenotypically wild-type *ssl/+* siblings unless indicated otherwise (**Figure 1B,C**). By mid-larval stages (7.7 mm standardized standard length, 7.7 SSL; ∼14 days post-fertilization), wild-type fish have newly differentiating adult melanophores dispersed over the flank as well as adult melanophores and persisting early larval melanophores along the horizontal myoseptum; iridophores have started to differentiate to form a primary interstripe and, in xanthophores, the first adult carotenoid pigmentation has become evident (Parichy et al., 2009; Patterson et al., 2014; Saunders et al., 2019). In the mutant, fewer adult melanophores were apparent at this stage, iridophores tended to be sparse and xanthophore pigment was not evident. During subsequent development, melanophores continued to be somewhat fewer than in wild-type and regions where xanthophores had differentiated in wild-type were markedly deficient for these cells in the mutant (e.g., 9.4 and 11.6 SSL in **Figure 1B**), albeit with variation among individuals, consistent with phenotypic variability of adults (**Figure 1C**; **Figure 1—figure supplement 1**).

### Defects in xanthophore proliferation and differentiation in *ssl* mutant

Given the especially severe reduction in xanthophore complement and the dependence of melanophore pattern on xanthophores (Parichy and Turner, 2003a) we focused on how the *ssl* mutation affects these cells. We first asked whether the xanthophore deficit reflects a failure of differentiation—in which case unpigmented xanthophores should persist in the *ssl* mutant, similar to a xanthophore defect in hypothyroid fish (McMenamin et al., 2014). To test this possibility, we examined the distribution of unpigmented xanthophores using the transcriptional reporter *mitfa*:Eos (Lister et al., 1999; Korzeniwsky et al., 2025), in which photoconvertible Eos fluorescent protein is driven by regulatory elements of *melanocyte inducing transcription factor a*. This orthologue of mammalian *MITF* is expressed strongly in melanophore and xanthophore lineages and at lower levels in other neural crest derivatives (Parichy et al., 2000; Eom et al., 2012; Saunders et al., 2019; Subkhankulova et al., 2023). The morphologies of Eos+ xanthophores (pigmented or unpigmented) allows them to be distinguished from unpigmented melanoblasts and earlier progenitors (Budi et al., 2011; Korzeniwsky et al., 2025) (**Figure 2—figure supplement 1A,B**).

**Figure 2.**
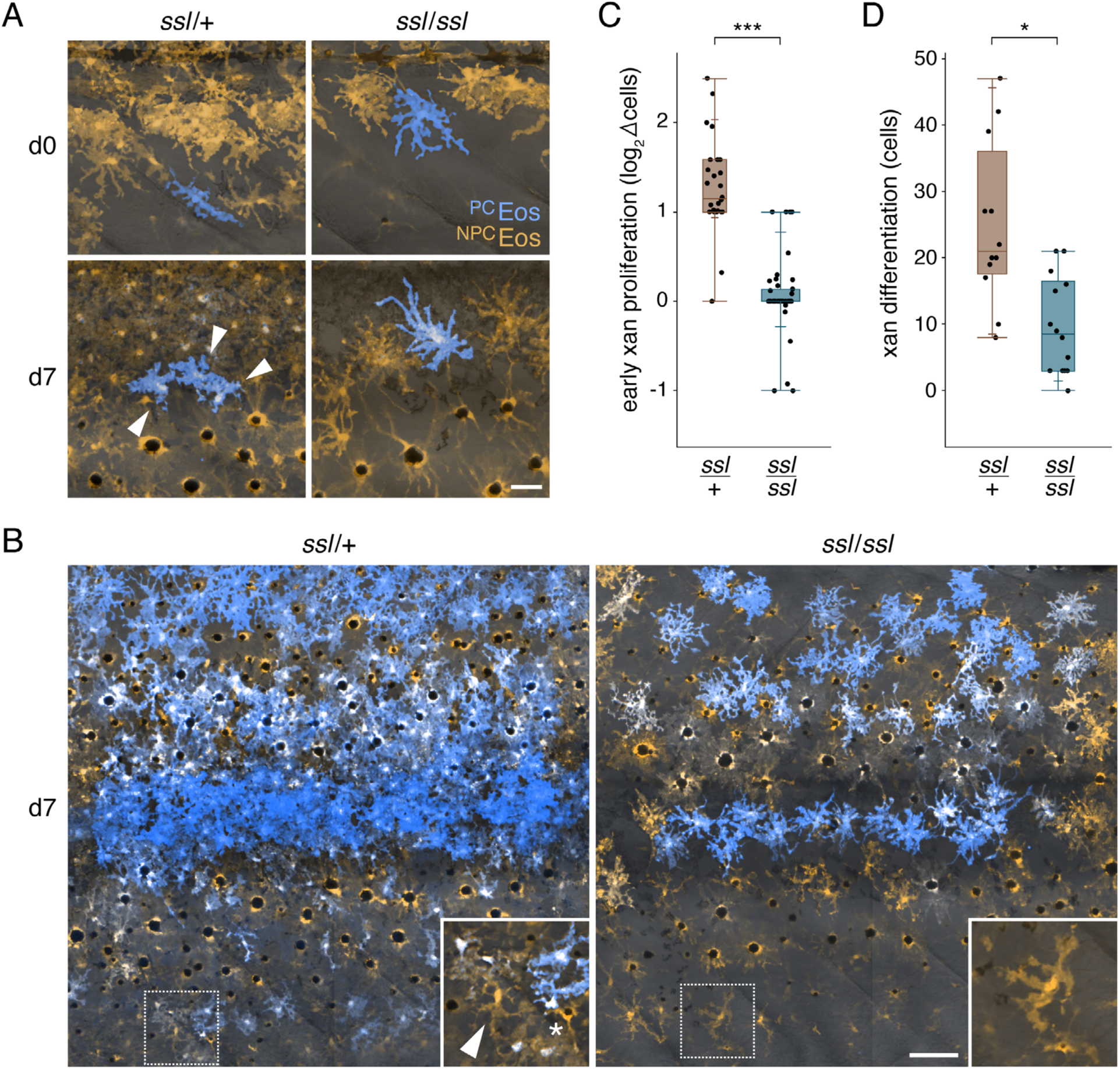
*ssl* mutant defects in xanthophore proliferation and differentiation. (**A,B**) Examples of wild-type and *ssl* mutant larvae in which cryptic xanthophores from embryonic stages were marked by Eos-photoconversion at ∼6.5 SSL (d0) and then re-examined at ∼8.6 SSL (d7). Photoconverted Eos (^PC^Eos) is shown in blue and non-photoconverted Eos (^NPC^Eos) is shown in orange. Continued transgene expression results in accumulation of non-photoconverted Eos on subsequent days, with overlap of converted and unconverted fluorophore appearing white. Here pseudocolored fluorescence images are shown superimposed on monochromatic brightfield images. (**A**) Individually photoconverted xanthophores of wild-type larvae divided to form small clones, in this case comprising three cells (arrowheads). By contrast, xanthophores of *ssl* mutants divided only rarely. Melanophores have melanin containing melanosomes concentrated towards cell centers by epinephrine treatment, with fluorophore evident in more peripheral regions. (**B**) After photoconverting cells over large regions, progeny of early xanthophores were evident dispersed widely over the flank, including ventral regions, where some newly differentiating xanthophores were also apparent. Inset shows a xanthophore without photoconverted fluorophore evident (arrowhead) surrounded by xanthophores with varying levels of perduring photoconverted fluorophore. A newly differentiating melanophore is marked with an asterisk. In *ssl*, early marked xanthophores mostly failed to increase in number or to colonize ventral regions of the flank, where xanthophores differentiating from latent progenitors appeared at times post-photoconversion (e.g., inset). (**C**). Proliferation of EL-derived xanthophores, here shown as doublings in cell number between d0–7, was markedly reduced in *ssl* compared to wild-type (Wilcoxon S=1594, *P*<0.0001, *n*=28 heterozygous and *n*=44 *ssl* mutant siblings). Plot boxes show median±IQR (interquartile range); whiskers extend to limits of data. (**D**) Numbers of xanthophores differentiating from latent progenitors, post-photoconversion of whole-fields, were also fewer in mutants compared to wild-type (Wilcoxon S=224, *P*=0.0015; *n*=12 ssl/+ and *n*=14 *ssl/ssl* siblings). Analyses of new xanthophore densities, normalizing for field-areas, yielded qualitatively similar results (*P*=0.0064). *, *P*<0.01; ***, *P*<0.0001. Scale bars, 50 µm (**A**), 100 µm (**B**).

**Figure 2—figure supplement 1.**
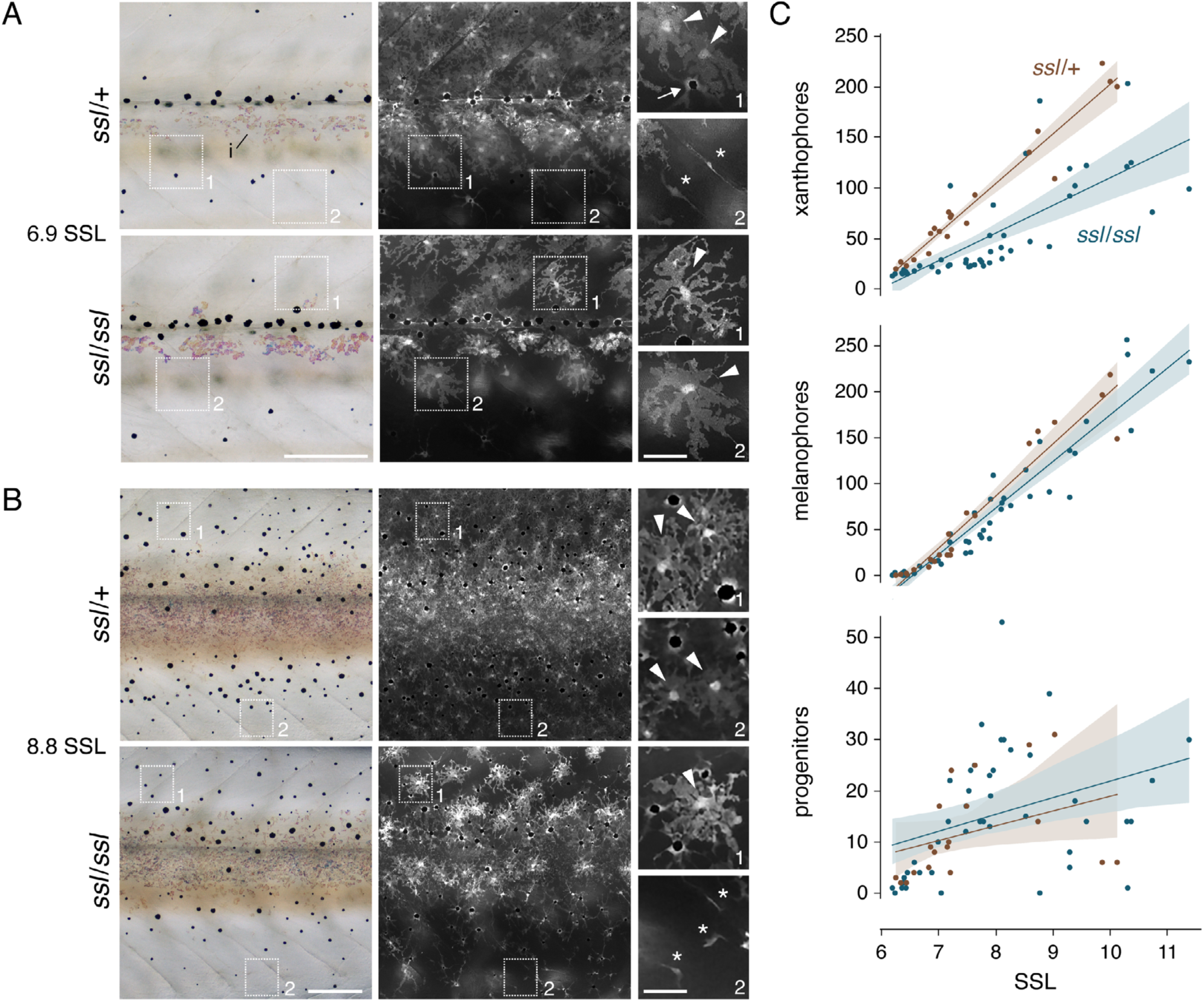
Complements of unpigmented *mitfa*:Eos+ cells in wild-type (*ssl/+*) and mutant (*ssl/ssl*) siblings. (**A**, **B**) Corresponding brightfield and fluorescence views at low magnification with details of boxed regions at higher magnification; two stages of development are shown (6.9 SSL; 8.8 SSL). Fewer unmelanized *mitfa*:Eos+ cells were evident in *ssl* homozygotes compared to phenotypically wild-type, heterozygous siblings. Arrowheads in details indicate individual Eos+ cells with morphologies characteristic of pigmented xanthophores, still-cryptic xanthophores derived from the early larval pigment pattern, or xanthoblasts derived from latent progenitors that have not yet differentiated with visible pigment (McMenamin et al., 2014; Eom et al., 2015; Korzeniwsky et al., 2025). Arrow in top panel indicates an Eos+ melanophore, with melanin at its center following epinephrine treatment. Asterisks mark smaller cells with spindle shapes typical of highly migratory earlier progenitors (Budi et al., 2011). i, accumulations of crystalline guanine reflecting platelets within iridophores of the developing interstripe (Patterson and Parichy, 2013; Gur et al., 2020). Display levels have been adjusted in detail panels to facilitate visualization. (**C**) After controlling for effects of size, numbers of *mitfa*:Eos+ xanthophores (cryptic or pigmented) differed significantly between genotypes (GLM Wald χ^2^=33.6, *P*<0.0001) whereas numbers of *mitfa*:Eos+ cells bearing melanin or having morphologies of progenitors were indistingushable between genotypes (both *P*>0.1) even after controlling for differences due to size or genotype x size interaction (*ssl/+*, *n*=20 fish; *ssl/ssl*, *n*=45 fish). Scale bars, 200 µm (overviews); 50 µm (details).

**Figure 2—figure supplement 2.**
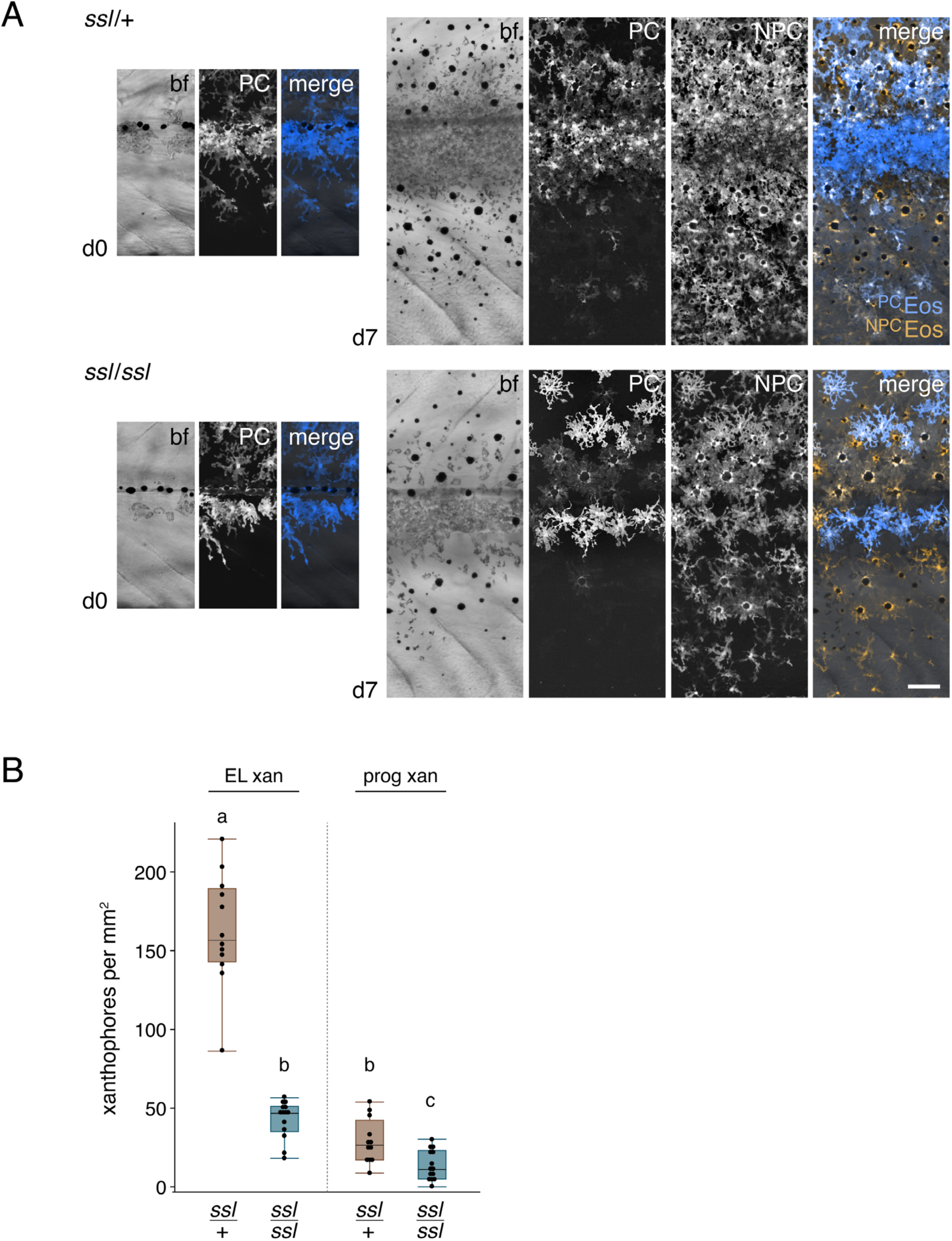
Photoconversions of wild-type and *ssl* mutant larvae. (**A**) Shown are same individuals as in Figure 2B on day of photoconversion (d0) and after one additional week of development (d7). Channel separations show monochrome brightfield (bf), photoconverted Eos (PC), non-photoconverted Eos (NPC), and merged (PC, blue; NPC, orange). (**B**) Contributions of EL-derived and progenitor-derived xanthophores to the total complement of this lineage, shown as cell density per mm^2^. Most xanthophores were EL-derived with lesser contributions in wild-type from progenitor-derived xanthophores; both of these populations were deficient in *ssl* mutants (overall ANOVA using *ln*-transformation to normalize variance of residuals, *F*3,48=17.9, *P*<0.0001; letters above bars indicates means not significantly different by post hoc Tukey-Kramer comparisons). Scale bar, 100 µm.

At stages spanning adult pigment pattern formation (4.4–11.6 SSL) *ssl* mutants had far fewer Eos+ xanthophores than wild-type siblings (**Figure 2—figure supplement 1C**) excluding a model in which *ssl* is required only for terminal differentiation. To determine if cells of the xanthophore lineage were not only reduced in number but also delayed or otherwise defective in differentiation, we compared the developmental stages / sizes (Parichy et al., 2009) at which pigmented xanthophores were first evident (**Figure 1C**). Differentiated xanthophores developed by 7.1±0.1 (mean±SE) SSL in wild-type, but these cells were first evident later, at 7.9±0.1 SSL, in the mutant (*n*=4 each, Wilcoxon S=26, *P*=0.03). *ssl* thus contributes to both normal numbers of xanthophore precursors and the timing of their differentiation.

Xanthophores of the adult have at least two origins. Many develop from yellow pteridine-containing xanthophores that first differentiate in the embryo to form the early larva (EL) pattern, evident between 3–6 dpf (Kimmel et al., 1995; Lister, 2019); as larvae grow, EL xanthophores divide and lose their pigments, assuming a cryptic state, then some redifferentiate with yellow-orange carotenoids in the light interstripes and others remain unpigmented or nearly so in the stripes (McMenamin et al., 2014; Saunders et al., 2019). Other xanthophores differentiate only during formation of the adult pigment pattern, arising from latent, neural crest derived progenitors (Parichy and Turner, 2003a; McMenamin et al., 2014).

To determine if *ssl* promotes the proliferation of EL-derived xanthophores, we marked individual Eos+ cryptic xanthophores by photoconversion and counted marked cells one week later (**Figure 2A**). In additional larvae, we photoconverted Eos+ cryptic xanthophores across a broad field (**Figure 2B**; **Figure 2—figure supplement 2A**) and assessed final numbers after the same interval. Estimates of proliferation were indistinguishable between approaches (method, *P*=0.6) so datasets were pooled for analysis. Numbers of EL-derived xanthophores in wild-type more than doubled during this period (median log2 fold-change, log2FC=1.15), whereas numbers in *ssl* mutants were virtually unchanged (median log2FC=0.00) (**Figure 2C**). *ssl* thus promotes the proliferation of cryptic xanthophores. Once marked, photoconverted cells only rarely disappeared in *ssl* mutants, arguing against a substantial role for cell death in contributing to xanthophore deficiency.

To learn if *ssl* also helps to expand the population of xanthophores from latent progenitors we compared numbers of newly arising xanthophores—lacking photoconverted fluorophore—between field-converted wild-type and mutant larvae (**Figure 2B**). Progenitor-derived xanthophores contributed only ∼15% of the total complement of xanthophores in phenotypically wild-type (*ssl/+*) fish, far less than EL-derived xanthophores (**Figure 2—figure supplement 2B**), and consistent with prior observations (McMenamin et al., 2014). As for EL-derived xanthophores, progenitor-derived xanthophores were significantly fewer in *ssl* mutants (**Figure 2D**), indicating that *ssl* promotes differentiation of latent progenitors towards a xanthophore fate, the proliferation of these cells, or both. Together, these analyses demonstrate roles for *ssl* in allowing normal numbers of xanthophores across distinct sublineages, with substantial defects in proliferation and differentiation evident in mutants.

Differentiated stripe melanophores very rarely divide (McMenamin et al., 2014) and corresponding analyses of field-photoconverted melanophores, at the same stages and over the same interval, did not reveal differences in melanophore proliferation or survival between wild-type and *ssl* mutant siblings (median±IQR log2FC=0.05±0.20, 0.00±0.20; *n*=16, 18 fish, respectively; Wilcoxon S=322, *P*=0.14).

Indeed, analyses of *mitfa*:Eos+ cells did not reveal differences between *ssl/+* and *ssl/ssl* in numbers of cells that contained melanin, or having a progenitor morphology (both *P*>0.1; **Figure 2—figure supplement 1C**). Rates of somatic growth and other conditions can impact pigment cell complements and pattern with some mutants showing substantial condition-dependence of phenotype (Parichy et al., 2003; Parichy et al., 2009): the modest deficiency in melanophores among individually reared *ssl/ssl* mutants (**Figure 1C**), not evident in group-reared fish, may reflect such a sensitivity and is consistent with variable expressivity of the phenotype more generally.

### *ssl* mutation dysregulates genes and pathways required for ribosome biogenesis and RNA processing

To identify genes and pathways that might explain defects in xanthophore development, we compared transcript abundances between *mitfa*:Eos+ cells of *ssl/+* and *ssl/ssl* siblings by single cell RNA-Sequencing (scRNA-Seq) at a late step in adult pattern formation, when larvae had reached the juvenile stage of 12 SSL (Parichy et al., 2009). Cells isolated by fluorescence-activated cell sorting included chromatophores and other neural crest derived cell types (e.g., progenitors, neurons and glia) that express low levels of the transgene (**Figure 3—figure supplement 1A–D**). We detected somewhat greater numbers of genes expressed in xanthophores and other cells of *ssl* mutants compared to wild-type (**Figure 3A**; **Figure 3—figure supplement 1E**). Preliminary analyses revealed significant differences in mutational effects across cell types (genotype x cell type interactions *q*<0.01 for 56% of loci tested), consistent with the differential impacts on cellular phenotypes we observed for xanthophores and melanophores. To understand potentially different responses of chromatophore classes we therefore subsetted the full dataset and compared *ssl* genotypes within cell types. scRNA-Seq recapitulated the xanthophore defects seen by fate mapping and repeated imaging. In *ssl* homozygotes, transcripts marking late steps of xanthophore differentiation [compiled from trajectory analyses of (Saunders et al., 2019)] were reduced relative to heterozygotes, and G1–S scores were elevated while G2–M scores were reduced, consistent with fewer xanthophore divisions (**Figure 3B**).

**Figure 3.**
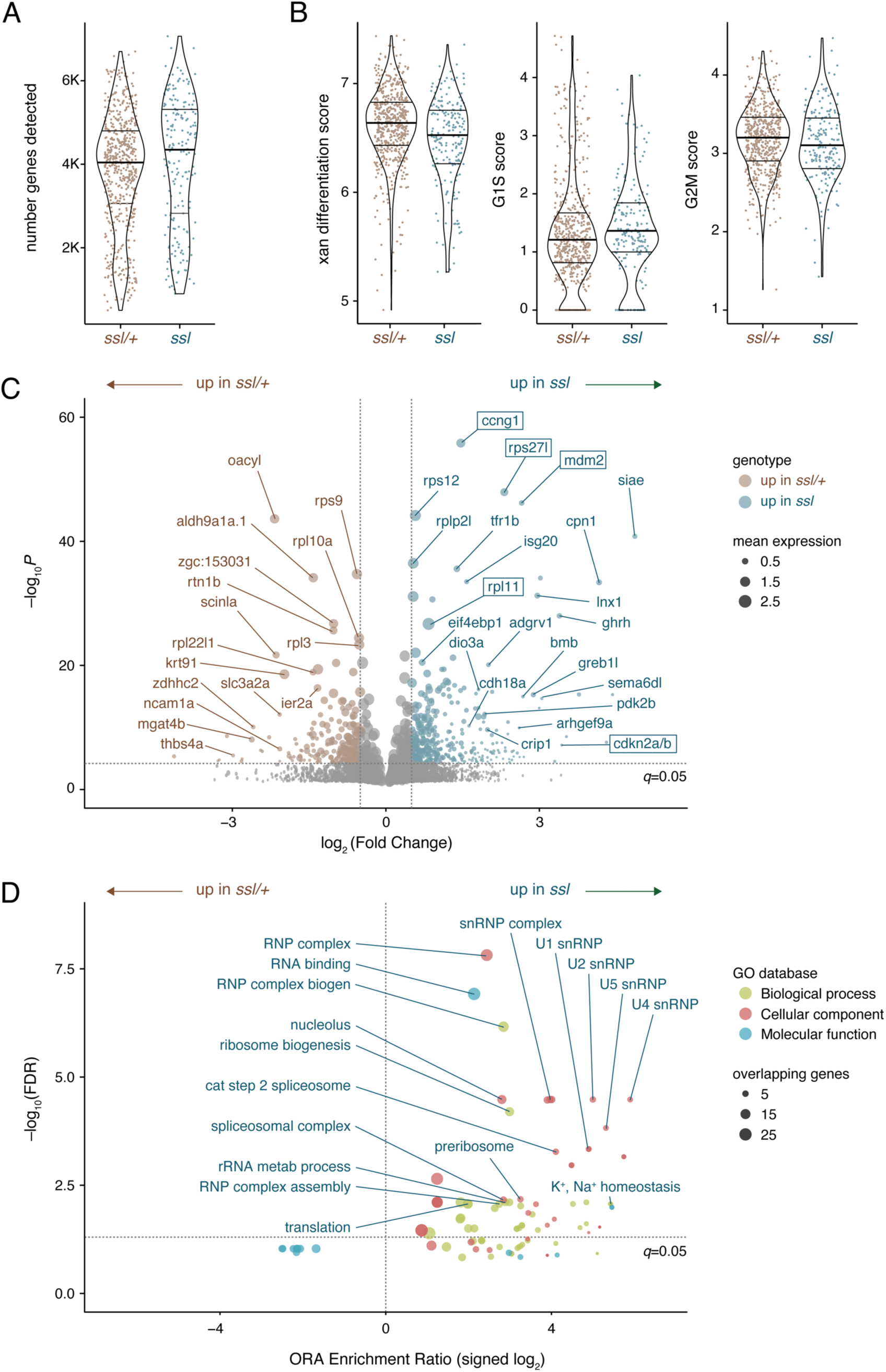
Molecular phenotype of xanthophore defect and pathways affected by *ssl* mutation. (**A**) Genes detected, showing values for individual cells and quartiles of data within violin plots (thick bars show medians). (**B**) Gene signature scores for xanthophores by scRNA-Seq (all signature score gene lists are in **Supplementary file 1—Table 1**). (**C**) Volcano plot illustrating gene expression differences in subsetted xanthophores (**Supplementary file 1—Table 2**). Point sizes are scaled to mean expression level within xanthophores of the genotype having higher expression levels. Several genes required for ribosome biogenesis or RNA processing were apparent amongst those expressed most differentially between genotypes. Of 18k genes detected in xanthophores, 377 were considered differentially expressed (*q*<0.05, |log2FC| > 0.5). (**D**) ORA from WebGestalt (Elizarraras et al., 2024) for GO terms with points colored for database and sized by numbers of genes identified as differentially expressed in xanthophores (ORA and GSEA results in **Supplementary file 1—Table 3, 4**).

**Figure 3—figure supplement 1.**
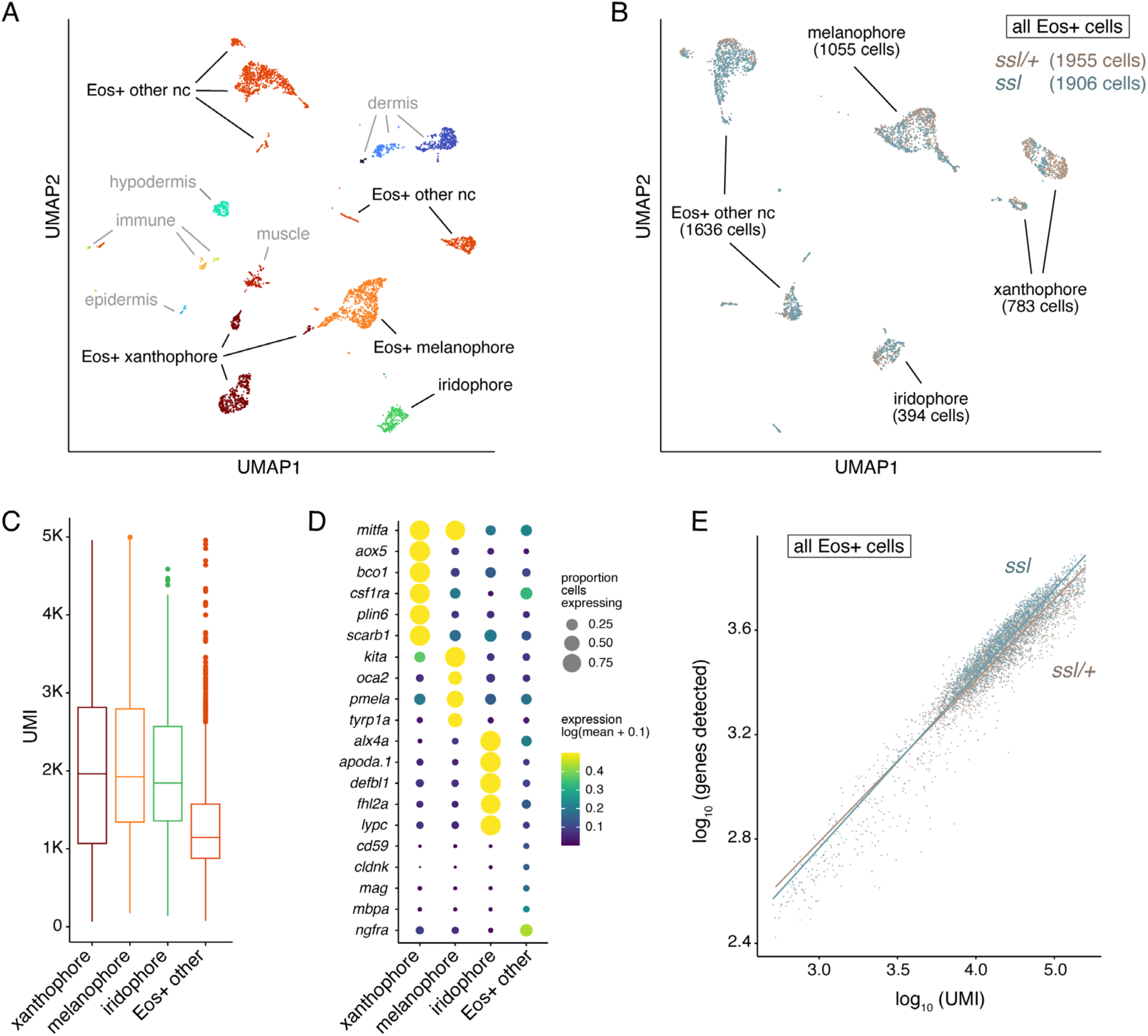
scRNA-seq from *ssl* mutants and wild-type siblings. (**A**) Unsupervised UMAP clustering and cell types manually assigned from analyses of differentially expressed marker loci. Broad cell-type categorizations are shown, sometimes encompassing more than a single cluster. Cell types known to express *mitfa:Eos* (Korzeniwsky et al., 2025) are indicated in black typeface; cell types representing presumptive by-catch are shown in grey. (**B**) UMAP representation of cells after removal of by-catch and reclustering, with cells per broad cluster or genotype indicated. Final datasets included 584 *ssl/+* xanthophores and 199 *ssl/ssl* xanthophores along with 555 *ssl/+* melanophores and 493 *ssl/ssl* melanophores. (**C**) Unique molecular identifiers (UMI) across cell types in B. (**D**) Expression of diagnostic marker genes for pigment cells, and other neural crest derived cells. (**E**) Numbers of genes detected were similar or greater in *ssl* mutant cells as compared to wild-type cells at a given depth of sequencing per cell (reflected in UMI).

**Figure 3—figure supplement 2.**
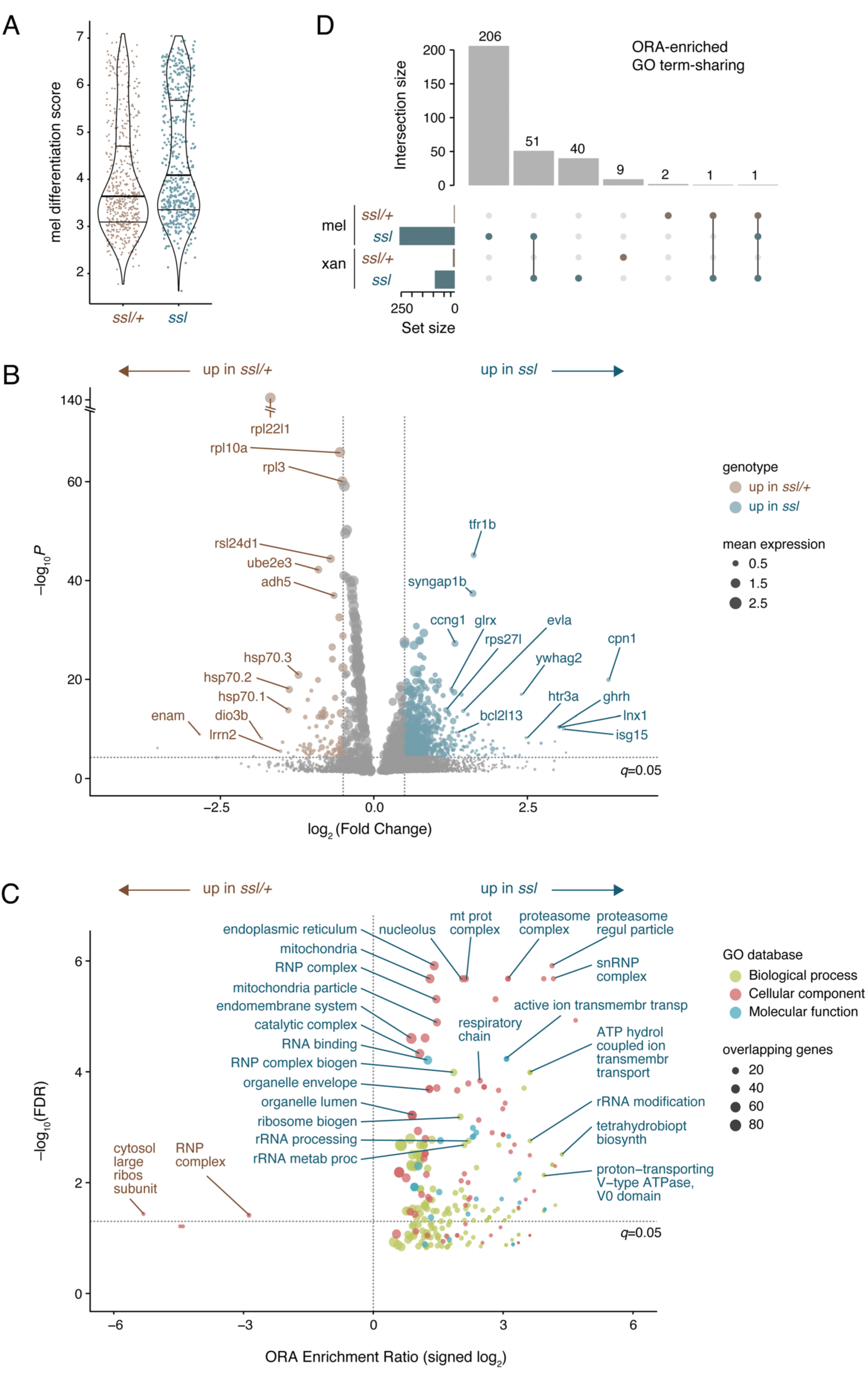
Gene expression changes in melanophores owing to *ssl* homozygosity. (**A**) Signature score for differentiation of melanophores. (**B**,**C**) Differentially expressed genes (*n*=689) (**B**) and overrepresented GO terms (**C**) in melanophores of *ssl/+* and *ssl/ssl*; lists for these and GSEA analyses can be found in **Supplementary file 1—Tables 2–4**. The larger number of differentially expressed genes for melanophores than xanthophores may reflect differences in statistical power given lesser sampling of the latter (Figure 3 **— figure supplement 1B**). (**D**) Upset plot (Lex et al., 2014) illustrates overlap for 51 (of 310) ORA-enriched GO terms between xanthophores and melanophores of mutants (shared and unique differentially expressed genes and GO term enrichment in **Supplementary file 1—Tables 5 and 6**, respectively).

**Figure 3—figure supplement 3.**
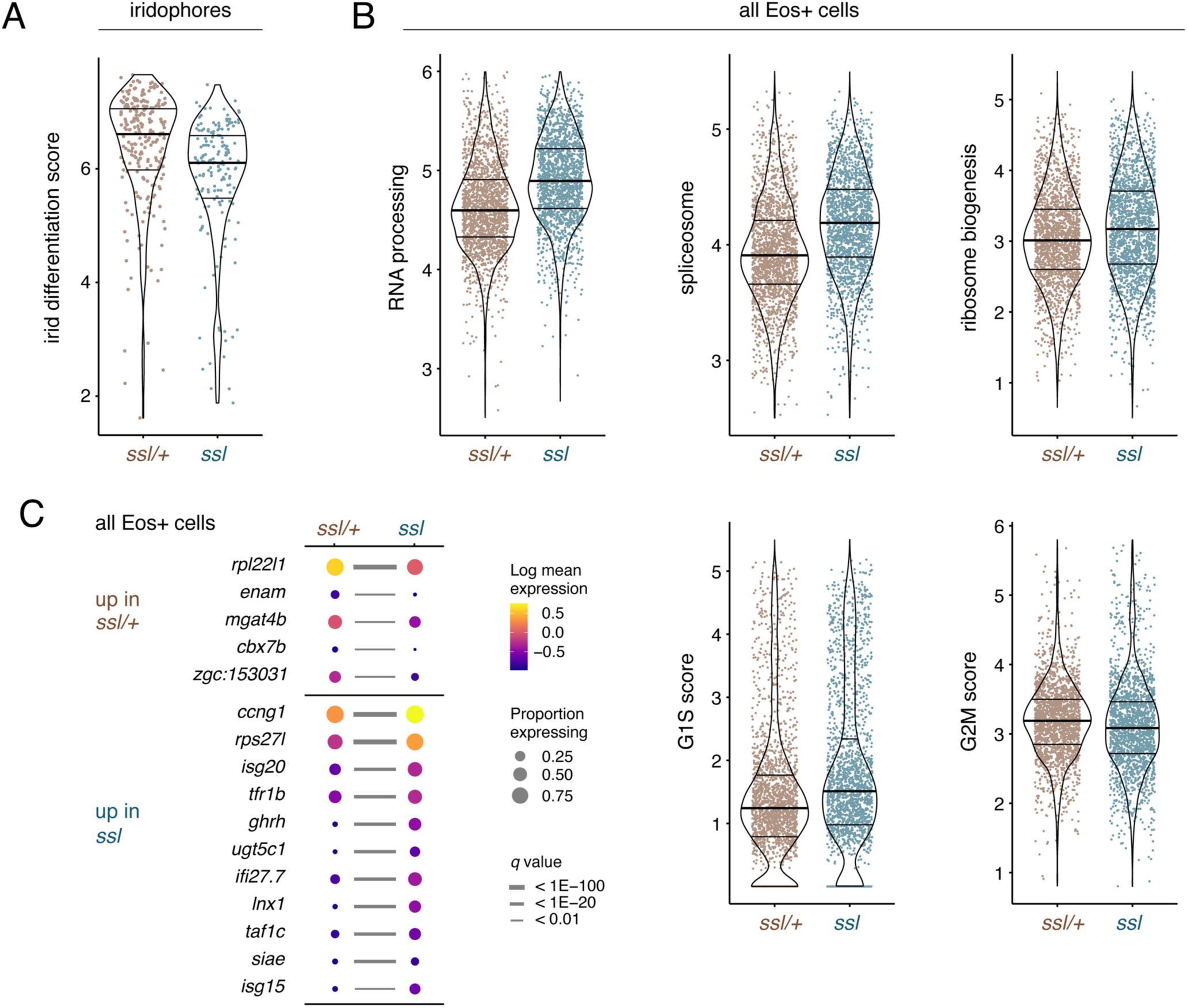
Transcriptomic analyses of iridophores and pooled *mitfa*:Eos+ cells. (**A**) Iridophore differentiation scores were reduced in *ssl* mutants though relatively few genes were differentially expressed between genotypes (*n*=103; **Supplementary file 1—Table 2**). (**B**) Signature scores across all *mitfa*:Eos+ cells extended inferences of dysregulated activities related to RNA processing, spliceosome components and ribosome biogenesis, along with modulated expression of cell cycle genes, particularly G1S markers. (**C**) Genes most differentially expressed between *ssl/+* and *ssl* mutants across all *mitfa*:Eos+ cells after controlling for cell-type specific differences.

Among the most differentially expressed genes were several encoding proteins of the small and large ribosomal subunits (**Figure 3C**). Also upregulated in mutant xanthophores, and distinct from core G1–S and G2–M markers, were several loci that couple ribosomal protein homeostasis to Tp53-dependent cell cycle arrest, including *mdm2*, *ccng1*, *rps27l*, *rpl11* and *cdkn2a/b* (see below).

To further investigate transcriptomic changes and consequences for pathway activities we used over-representation analyses (ORA) for gene ontology (GO) terms of biological processes, cellular components and molecular functions (**Figure 3D**). These revealed in *ssl* mutants an upregulation of genes associated with ribosomes, ribosome biogenesis, ribonucleoprotein complexes, nucleoli and translation, as well as RNA processing and splicing. No GO categories were overrepresented among genes more highly expressed in *ssl/+*. Gene set enrichment analyses (GSEA) yielded a concordant outcome. Together these findings suggested generalized defects related to ribosomes and RNA processing, rather than alterations in regulatory programs specific to xanthophore differentiation or morphogenesis.

To learn if qualitatively similar impacts of *ssl* mutation extend beyond xanthophores, we examined differentially expressed genes of melanophores and other cells. Surprisingly, a differentiation signature score for melanophores was elevated in homozygous mutants, perhaps reflecting lower density and greater availability of differentiation-promoting factors, or a reduced number of cells at early states of specification (**Figure 3—figure supplement 2A**). Differential expression analyses, ORA and GSEA revealed similar trends as in xanthophores, namely large effects on genes having ribosome-associated GO terms, with little indication of melanophore-lineage specific response; terms associated with spliceosome complexes were over-represented though not as prominently as in xanthophores (**Figure 3—figure supplement 2B,C**). About 17% of ORA-enriched GO terms (51 of 310) were shared between melanophores and xanthophores of mutants, representing ribosome biogenesis, RNA processing, and translation as well as monovalent cation homeostasis (e.g., Na⁺/K⁺ and ion-transporting ATPase activity and homeostasis), consistent with roles for bioelectric signaling in development of these cell types (Iwashita et al., 2006; Inaba et al., 2012; Klann et al., 2026)(**Figure 3—figure supplement 2D**).

By contrast with melanophores, iridophores were substantially reduced in differentiation signature score in *ssl/ssl* compared to *ssl/+* though only comparatively few genes were differentially expressed (**Figure 3—figure supplement 3A**); no GO terms sets were found to be differentially enriched for these cells.

We next considered whether trends suggested by differentially expressed genes and GO enrichments were true of *mitfa*:Eos+ cells more generally. As compared to wild-type cells, mutant cells had higher signature scores for RNA processing, spliceosome, ribosome biogenesis and G1–S scores and a marginally reduced G2–M score (**Figure 3—figure supplement 3B**). Finally, we asked whether any genes were especially affected by the *ssl* mutation across the entire dataset of *mitfa*:Eos+ cells and so generated contrasts to isolate shared mutational effects across all cell-types, normalized for differential cell-type abundance and isolated from effects specific to cell-type classifications (i.e., xanthophore, melanophore, iridophore, other). Compared to cells of the wild-type, this analysis highlighted among other genes an upregulation in mutants of *ccng1* (Cyclin G1), *rps27l*, and *ghrh* (Growth hormone releasing hormone) and a concomitant downregulation of *rpl22l1* (**Figure 3—figure supplement 3C**).

### *ssl* is a mutant allele of DEAD-helicase gene *ddx49*, which functions autonomously in melanophores and xanthophores

Transcriptomic analyses pointed to roles for *ssl* in ribosome and RNA biology, functions likely to be manifested as autonomous effects of the gene on chromatophores, though interactions between chromatophores also can have substantial effects on differentiation and patterning of these cells (Parichy and Turner, 2003; Frohnhofer et al., 2013; Patterson and Parichy, 2013; Aman et al., 2023). Consistent with the cell-autonomy of *ssl* activity and narrowing the spectrum of potential candidate genes, wild-type donor cells transplanted to mutant hosts formed pigment pattern elements more similar to wild-type than *ssl* mutants (**Figure 4—figure supplement 1**).

**Figure 4.**
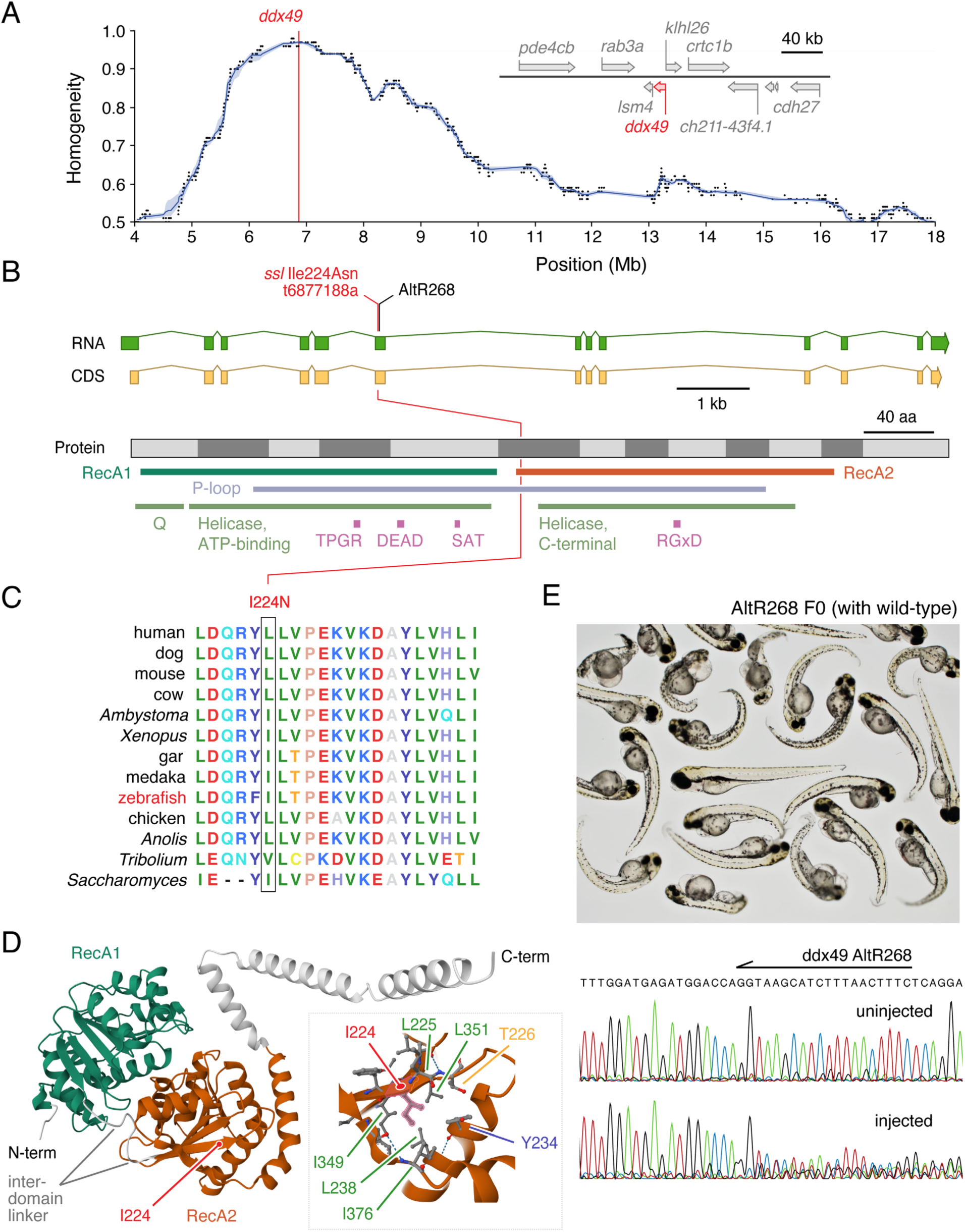
Mutation in DEAD-box helicase gene *ddx49* identified in *ssl* pigment pattern mutant zebrafish. (A) Whole-genome resequencing of pooled mutant DNA revealed a peak in homogeneity from ∼6–7.5 Mb on Chr11, a region containing *ddx49*, at upper right with neighboring genes between 6.72–7.04 Mb. (B) The region of high homogeneity included only one mutation predicted to be deleterious, leading to a missense substitution of Ile to Asn at position 224 of Ddx49. Conserved sequence features include: Helicase superfamily 1/2 ATP-binding domain; Helicase C-terminal domain-like; P-loop, P-loop containing nucleoside triphosphate hydrolase; Q, DEAD-box type RNA helicase Q motif; and DEAD-box. Ensembl ENSDARP00000015246.7, UniProt Q7ZUT9. RecA1 and RecA2 domains correspond to AlphaFold structure domains of the P-loop NTPase superfamily (CATH: 3.40.50.300) and are shown as colored regions of protein structure in D. (**C**) Multiple sequence alignment including I224 of zebrafish reveals conserved nonpolar, hydrophobic amino acids with branched side chains at the homologous positions across species. The homologous sequence shown from *Saccharomyces* corresponds to protein Dbp8p. (**D**) AlphaFold structure prediction for zebrafish Ddx49 (AF-Q7ZUT9-F1-v6) with context of I224N substitution shown in detail (slightly rotated to show other nonpolar hydrophobic amino acids nearby); dotted lines, predicted hydrogen bonds formed by backbone groups. (**E**) CRISPR/Cas9 targeting of zebrafish *ddx49* with AltR268 (site shown in B) led to embryonic lethal or larval-inviable phenotypes in genetically mosaic F0 fish, here surrounding an uninjected sibling control. Lower part of panel shows AltR268 target sequence and representative electropherograms from pooled uninjected controls and pooled injected embryos, indicating efficacious induction of somatic mutations [total efficiency of induced mutations=71.6% (Brinkman et al., 2014)].

**Figure 4—figure supplement 1.**
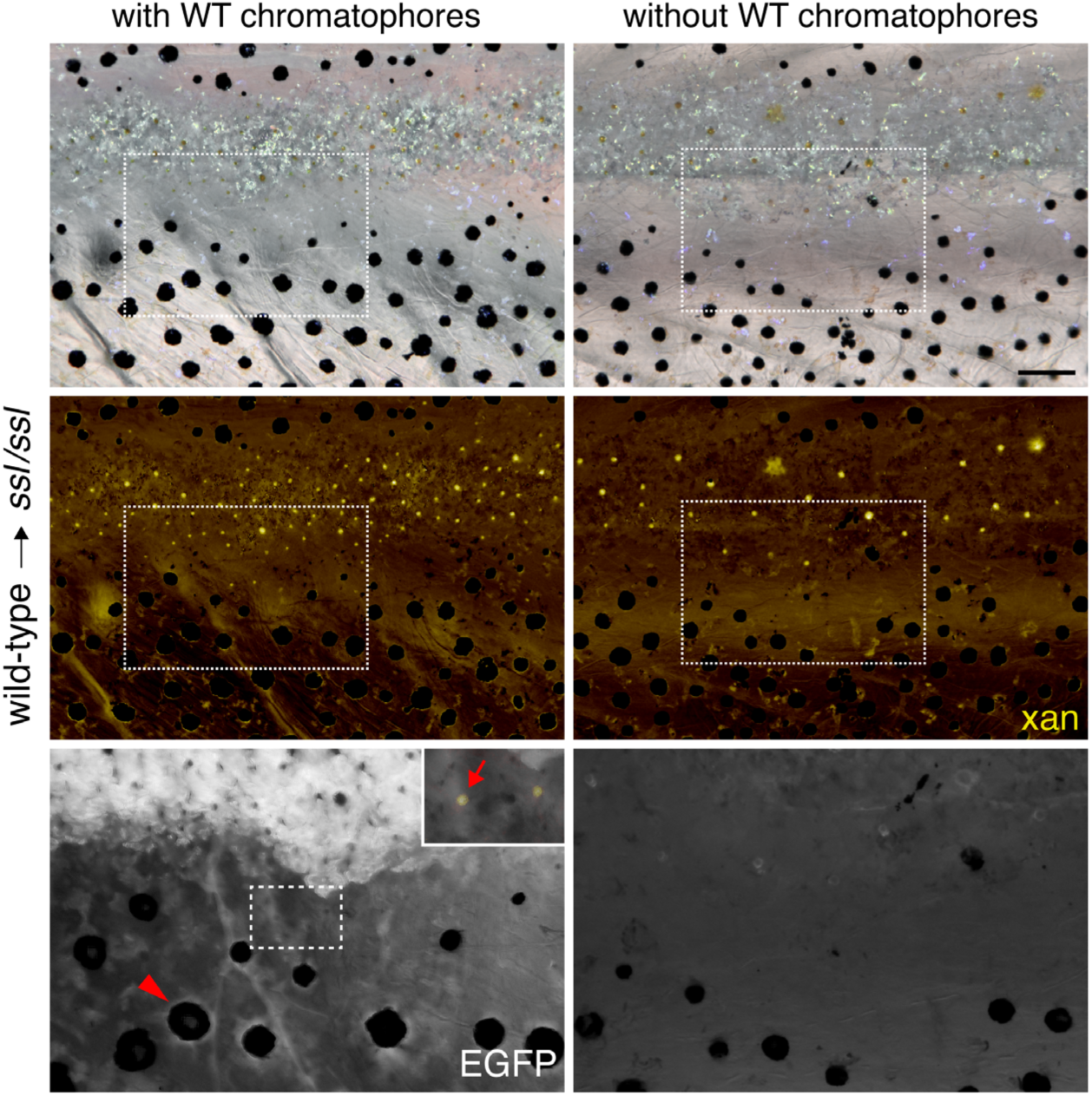
Blastula stage transplants suggest pigment cell autonomous *ssl* activity. Example of a chimeric fish in which wild-type cells, transgenic for widely expressed EGFP [*Tg(ubi:Switch)*] (Lalonde et al., 2022), were transplanted to an *ssl/ssl* mutant host. Left panels show a region where donor cells differentiated as pigment cells, with a regular pattern of stripes and interstripe. The interstripe has abundant EGFP+ xanthophores (brightfield pseudocolor in middle panel) with EGFP+ iridophores, and EGFP+ melanophores (arrow); xanthophores apart from iridophores are shown in bottom panel inset, illustrating pseudocolored carotenoid deposits. Right panels show a nearby region of the same individual, not colonized by donor wild-type chromatophores, where mutant xanthophores were sparse and patterning irregular. Overall, donor chromatophores developed similar patterns in each of 16 chimeras in which donor cells persisted into the adult; in 43 additional chimeras examined as adults, donor cells differentiated as epidermis, muscle, scales, nerves, and other cell types and changes in mutant pigmentation were not discernible. Scale bar, 100 µm.

**Figure 4—figure supplement 2.**
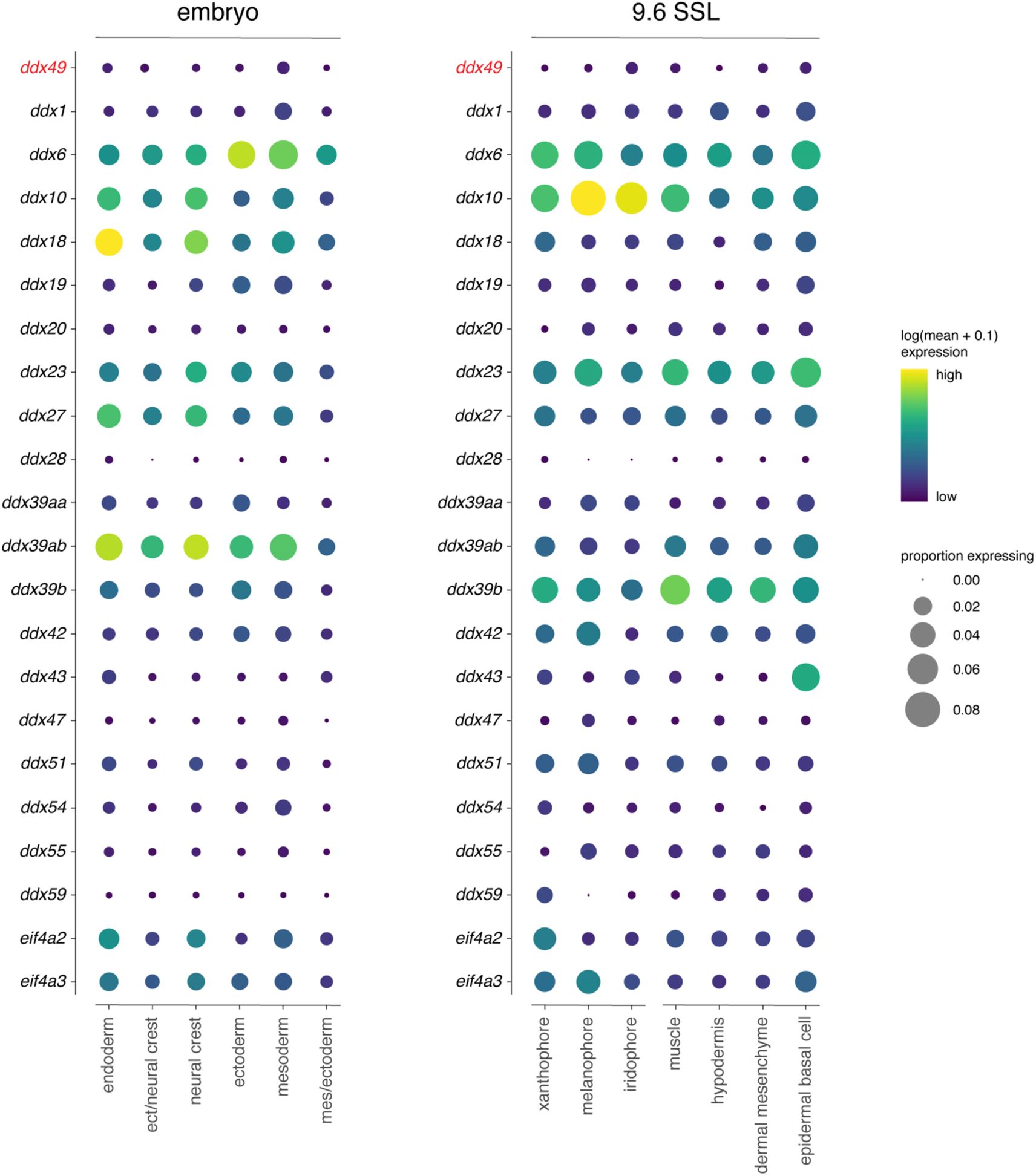
*ddx49* expression during embryonic development and larva-to-adult transformation of wild-type zebrafish. Dot plots indicate transcript abundance and proportion of sampled cells expressing *ddx49* from whole embryos (18–96 hours post-fertilization; 630k cells) (Saunders et al., 2023) or within the skin during late adult pattern development (9.2 SSL), including major pigment cell classes and other cells with which pigment cells may interact (10k cells total across types indicated) (Aman et al., 2023). Additional helicase genes are shown for comparison, identified as orthologues of human genes having protein products with the highest—albeit still low—sequence identities to human DDX49 (here 32–46%), or similarities of inferred protein interaction networks (also very limited) (Awasthi et al., 2018).

To identify the gene affected in *ssl* we crossed a homozygous mutant fish to the inbred wild-type strain WIK and then backcrossed a female F1 to *ssl/ssl*. From the resulting progeny we sequenced genomic DNA of pooled homozygous mutant and pooled heterozygous phenotypically wild-type fish. Subsequent analyses revealed a broad region of allelic homogeneity on Chr11 associated with *ssl* homozygosity, centered at ∼6.85 Mb (**Figure 4A**). Assessment of novel variants and likely effects between 4.5–11.0 Mb failed to reveal nonsense or canonical splice site mutations and only a single missense substitution in mutants, confirmed by Sanger sequencing, near the peak of homogeneity at 6.88 Mb. This variant occurred within exon 6 of *ddx49*, encoding a DEAD-box protein of the superfamily 2 (SF2) RNA helicases, having potential functions consistent with cell-autonomous effects (**Figure 4B**). The affected amino acid, Ile224, is conserved to be nonpolar hydrophobic across vertebrates, insects and yeast (**Figure 4C**) whereas the substitution, Asn224, is polar hydrophilic and predicted to be deleterious for zebrafish and likely pathogenic for the corresponding amino acid (Leu224) of human (SIFT score=0.00, AlphaMissense Pathogenicity score=0.944, respectively)(Ng and Henikoff, 2003; Tordai et al., 2024). The affected site is inward facing and just inside of the conserved C-terminal helicase (RecA2) domain (**Figure 4D**). scRNA-Seq indicated marginally more abundant *ddx49* transcript in *mitfa*:Eos+ xanthophores and all *mitfa*:Eos+ cells of *ssl* mutants than *ssl/+* siblings (log2FC=0.62 and 0.33, respectively; both *P*<0.002), consistent with feedback leading to a compensatory response in expression.

DEAD-box (DDX) proteins often localize to membraneless organelles, including processing bodies, stress granules, nuclear speckles and nucleoli, and have diverse, cell-autonomous functions in transcription, mRNA splicing and alternative splicing, mRNA transport, storage, and decay, regulation of translation, ribosome biogenesis, and innate immunity (Bohnsack et al., 2023; Khreiss et al., 2023; Dorner and Hondele, 2024; Tapescu and Cherry, 2024). Many DDX proteins are well-studied: e.g., the translation initiation complex factor eIF4A1/DDX2A, exon junction complex protein eIF4A3/DDX48, and the germ line specification factor VASA/DDX4. DDX49 and its yeast orthologue Dbp8 have received far less attention but, as noted above, have been implicated in the processing of pre-ribosomal RNAs (Daugeron and Linder, 2001; Awasthi et al., 2018). Such a role would be consistent with pathways affected by *ssl* mutation (**Figure 3C**) as well as widespread, albeit low expression of *ddx49* in embryos and post-embryonic skin, evident in reanalyses of previously published scRNA-seq data (**Figure 4— figure supplement 2**).

Given this potential function and apparent ubiquity of expression, we considered the possibility that null alleles might be embryonic-lethal. Indeed, targeting of a site near Ile224 to induce premature termination alleles and other variants by CRISPR/Cas9 mutagenesis (Hoshijima et al., 2019) resulted in mosaic embryos having a variety of morphological defects that did not survive beyond early larval stages (**Figure 4E**). Thus, *ddx49* is essential for early zebrafish development.

Early lethality of induced *ddx49* mutations made it impractical to confirm allelism with *ssl* by phenocopy or complementation testing with new alleles. We therefore tested by transgenesis if *ssl* mutant phenotypes result from dysfunction of Ddx49 I224N by injecting one-cell *ssl* mutant embryos with plasmid (*mitfa:nucVenus-2a-ddx49*) to express wild-type Ddx49 mosaically in xanthophores, melanophores and their progenitors, which resulted in partial or even seemingly complete rescues of pigment cell complements and patterns both on the body and in the fin (**Figure 5**; **Figure 5—figure supplement 1A,B**). Close inspection of these phenotypes further suggested autonomous effects within xanthophores and melanophores or their *mitfa*-expressing progenitors. Where *ssl* mutant xanthophores expressed wild-type *mitfa*:Ddx49, these cells tended to be widespread, extending beyond regions with *mitfa*:Ddx49+ melanophores, consistent with recovery of the substantial proliferative abilities of wild-type xanthophores (Parichy and Turner, 2003a; Walderich et al., 2016) (**Figure 5—figure supplement 1B**). Near xanthophores expressing wild-type *mitfa*:Ddx49, melanophores occurred in well-formed stripes whether or not they expressed the transgene; this outcome was not surprising given roles for xanthophores in promoting melanophore organization into stripes (Parichy and Turner, 2003a); likewise, although iridophores did not express the nucVenus reporter at discernible levels, we cannot formally exclude the possibility that functionally significant levels of *mitfa*:Ddx49 activity in these cells also contributed to melanophore organization. Where melanophores expressed *mitfa*:Ddx49, however, these cells had significantly greater densities (by ∼66±12%, mean±SE) than adjacent regions where melanophores lacked the transgene (matched pairs, *t*4=5.35, one-tailed *P*=0.003), consistent with melanophore-lineage autonomy of Ddx49 in promoting population expansion. Together these outcomes verified—for this late-appearing pigmentary phenotype—correspondence of *ssl* and *ddx49*, and implied that Ddx49 functions autonomously to expand populations of both xanthophores and melanophores, with non-autonomous effects of xanthophores on melanophores required for pattern organization.

**Figure 5.**
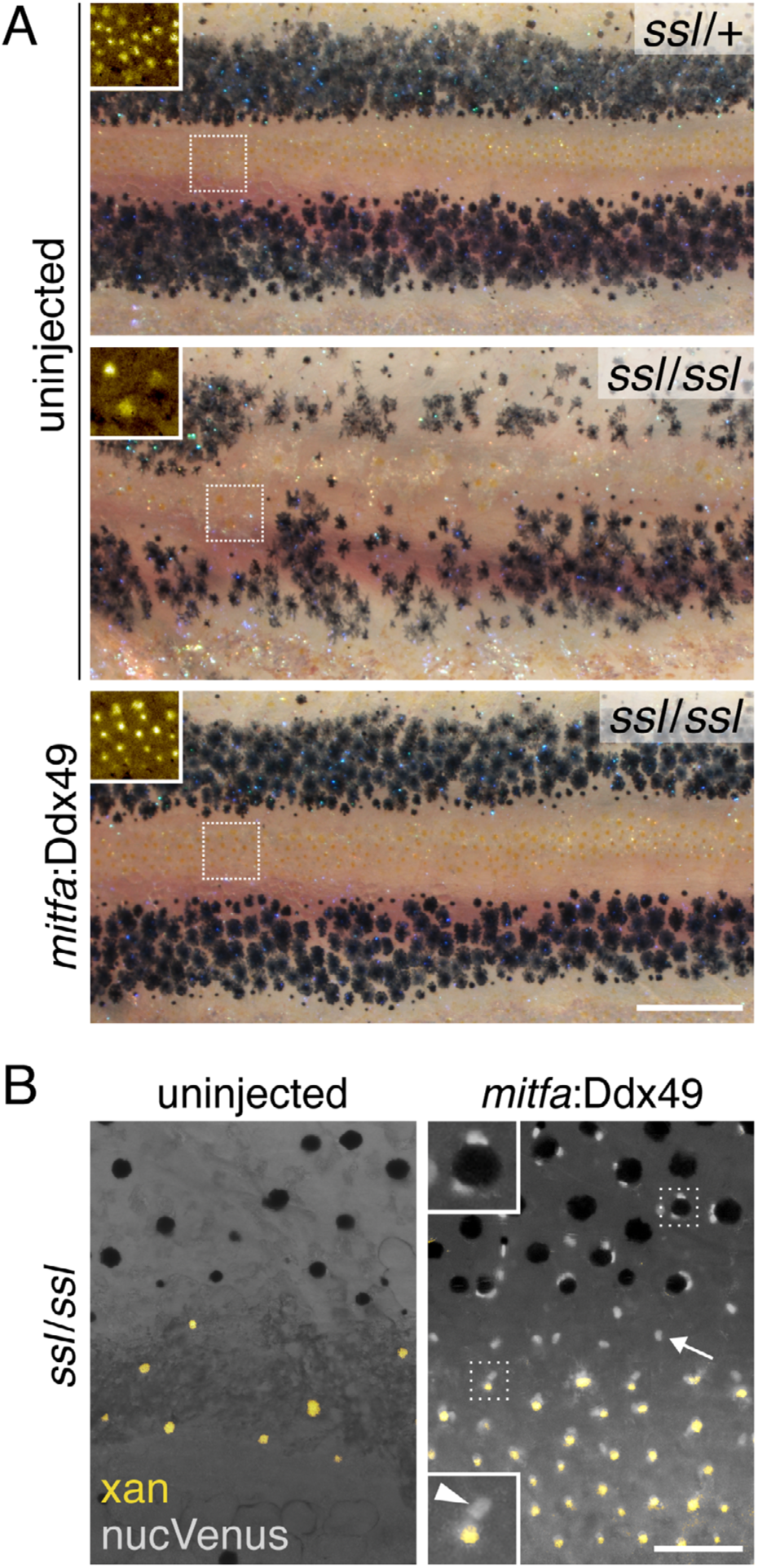
Ddx49 rescued *ssl* mutant phenotype. (**A**) Uninjected wild-type and mutant controls with representative mutant expressing wild-type Ddx49 in melanophores, xanthophores and their progenitors by injection of *mitfa:nucVenus-2a-ddx49* transgene. Inset squares show pseudocolored xanthophores. Similar outcomes were evident in each of 9 *mitfa*:Ddx49 expressing *ssl* mutant individuals examined at young adult stages. (**B**) Xanthophore pigment in pseudocolor (yellow) with nuclear localizing Venus fluorophore, linked to Ddx49 expression by 2a self-cleaving peptide sequence (Provost et al., 2007). Lower inset, oblong nucVenus+ nucleus (arrowhead) of xanthophore next to carotenoid vesicles (yellow) contracted towards cell centers (Huang et al., 2021). Upper inset, nucVenus+ nuclei adjacent to contracted melanosomes of melanophore, here binucleated as is typical of these cells (Saunders et al., 2019). Additional nucVenus+ nuclei were evident without associated pigment (e.g., arrow) reflecting expression of *mitfa* transgene in progenitors and cryptic xanthophores (Korzeniwsky et al., 2025). Scale bars: 500 µm (**A**), 100 µm (**B**).

**Figure 5—figure supplement 1.**
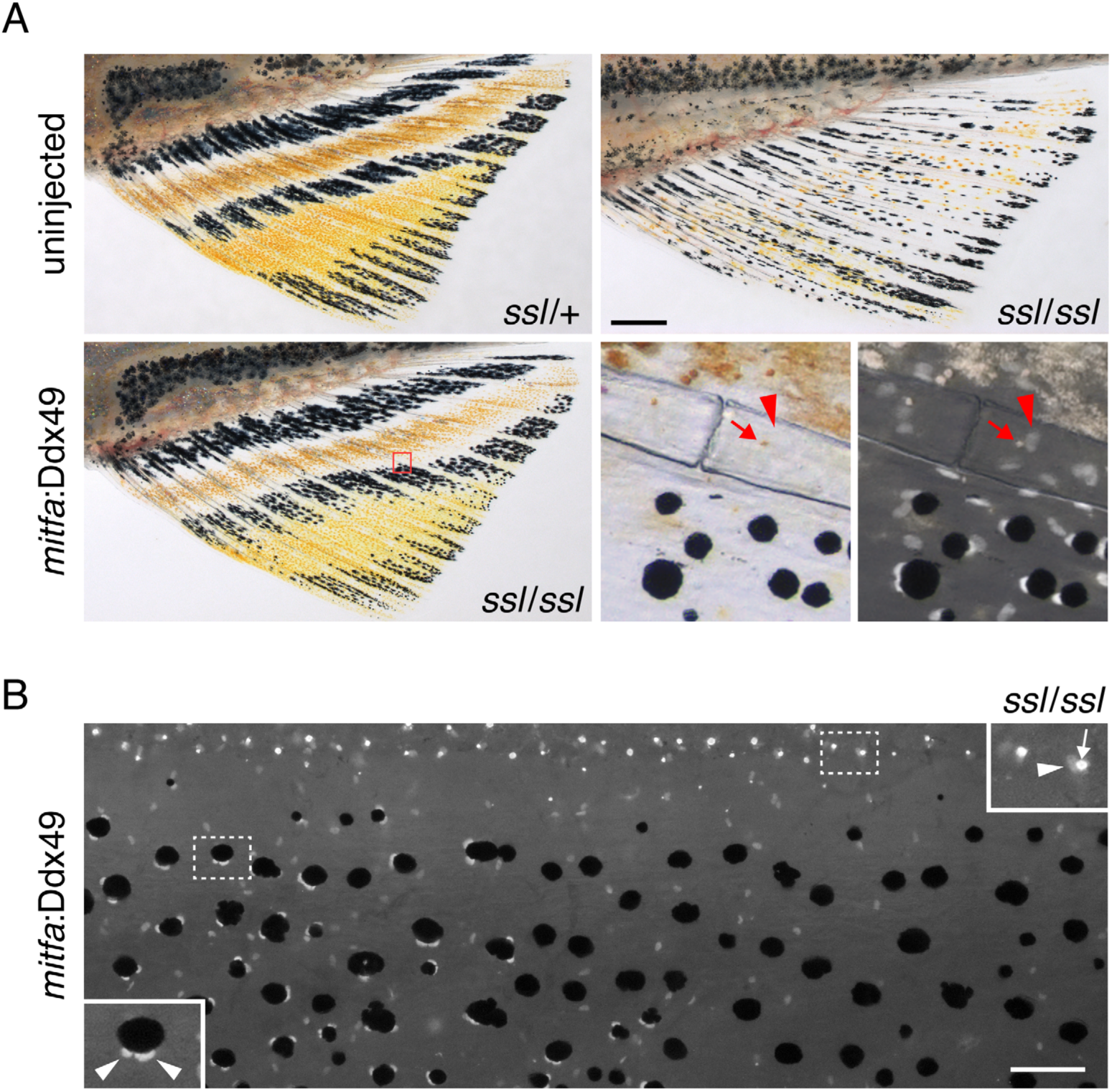
Recovery of pigment cell complements and pattern in *ssl* mutants. (**A**) Transgenic expression of *mitfa*:Ddx49 rescued melanophores, xanthophores and the stripe pattern in fins. Lower right panels correspond to region outlined by red box at left, following epinephrine treatment. Arrowhead, location of nucVenus+ xanthophore nucleus adjacent to autofluorescent carotenoid (arrow). Transgene-labeled nuclei of melanophores and unpigmented, presumptive progenitors are also evident at right. (**B**) *mitfa*:Ddx49 rescued xanthophores and interstripe/stripe pattern, shown here with transgene-expressing interstripe xanthophores across the top of the frame. Upper inset shows a nucVenus+ nucleus (arrowhead) adjacent to epinephrine-concentrated autofluorescing carotenoid vesicles in the same xanthophore (arrow). Where *mitfa*:Ddx49+ xanthophores were present, melanophores were arranged in stripes whether or not they expressed the transgene, as illustrated here with *mitfa*:Ddx49+ melanophores to the left (lower inset: arrowheads, nucVenus+ nuclei) and *mitfa*:Ddx49– melanophores to the right. Where melanophores expressed *mitfa*:Ddx49, densities were greater in comparison to adjacent regions where melanophores lacked the transgene. Scale bars, 500 µm (**A**), 100 µm (**B**).

### Dysregulation of Ddx49 activities by *ssl* mutation

We next sought to understand how the *ssl* Ile224Asn (I224N) substitution affects Ddx49 function and so assessed consequences for protein localization, abundance and activity. When expressed as GFP-tagged fusion proteins in human 293T cells both wild-type and I224N mutant proteins localized to nucleoli, as previously reported for human DDX49 (Awasthi et al., 2018) (**Figure 6A**). To assess Ddx49 stability and activities *in vitro* we overexpressed His-tagged wild-type and I224N Ddx49 in bacteria and affinity purified these proteins, as well as GST-tagged human DDX49 for use as a positive control (**Figure 6 — figure supplement 1A**). The I224N substitution did not induce major changes in protein secondary structure compared to wild-type, as assayed by circular dichroism spectroscopy (**Figure 6— figure supplement 1B**). Nevertheless, abundance of I224N Ddx49 was reduced compared to wild-type Ddx49, as revealed by quantitative comparisons of immunoblots, using coexpressed GFP-tagged GAPDH for normalization (**Figure 6B**; **Figure 6—figure supplement 1C**), suggesting reduced stability at least in this heterologous context.

**Figure 6.**
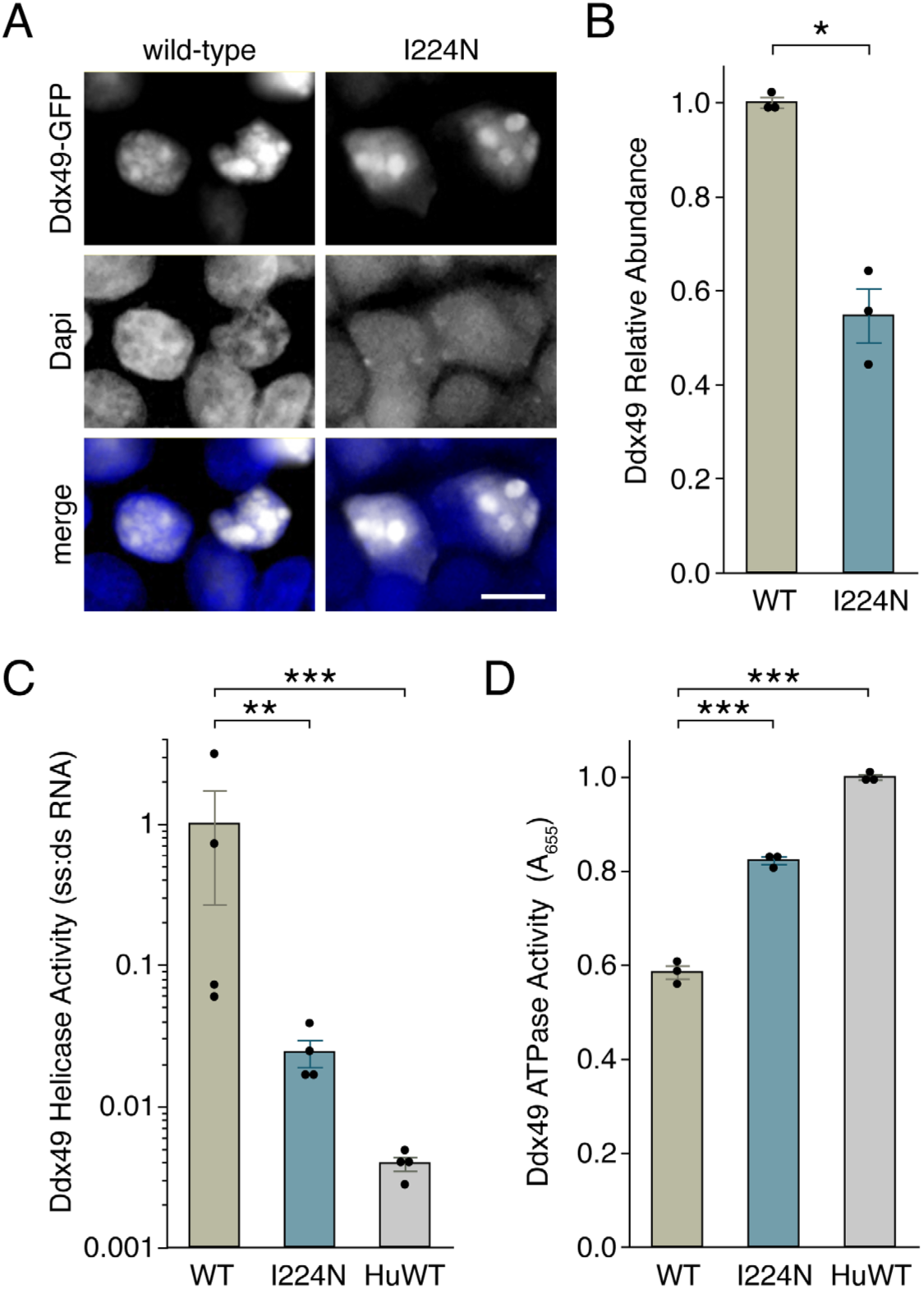
Biochemical assay of *ssl* mutant Ddx49 activities and stability. (**A**) Wild-type and I224N mutant GFP-Ddx49 both localized to nucleoli in human 293T cells. (**B**) Lesser abundance of Ddx49 I224N compared to wild-type (normalized to co-expressed GFP-GAPDH; *F*1,4=59.6, *P*<0.01). (**C**) Ddx49-I224N had greater ATPase activity than wild-type Ddx49, though human DDX49 (HuWT) had greater activity compared to either zebrafish protein (*F*2,6=452.9, *P*<0.0001). (**D**) Ddx49-I224N was reduced in helicase activity compared to wild-type Ddx49, though both forms showed considerably greater activity than human DDX49 (generalized linear model: *F*2,9=40.4, *P*<0.0001 with response=ssRNA:dsRNA, distribution=gamma, link function=log). Asterisks in (**C**) and (**D**) indicate *post hoc* Tukey-Kramer comparisons of means. \**P*<0.01; **, *P*<0.001; ***, *P*<0.0001. Scale bar,10 µm (**A**).

**Figure 6—figure supplement 1.**
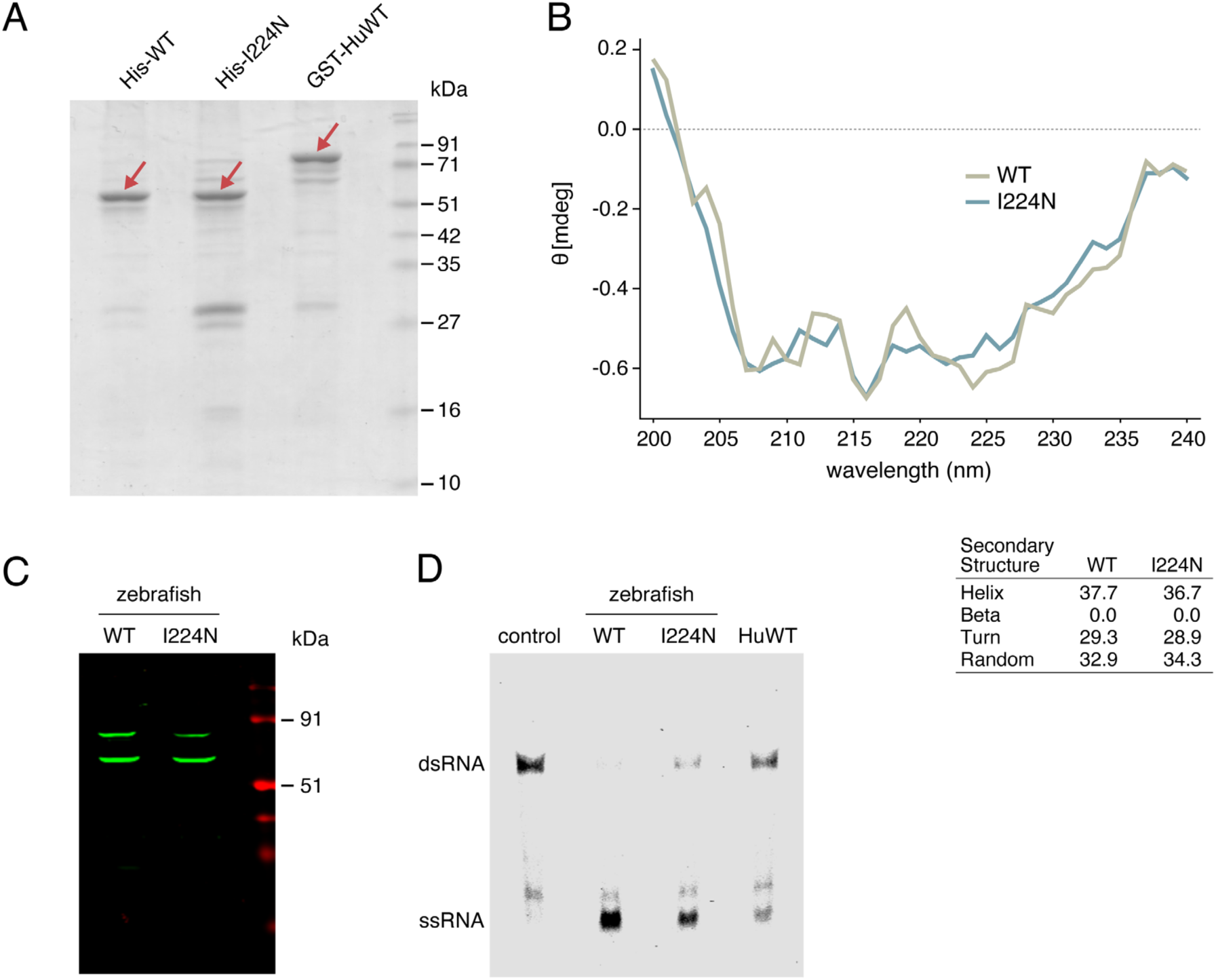
Mutant Ddx49 structure, expression and helicase activity. (**A**) Expression of His-tagged and GST-tagged proteins for *in vitro* assay. (**B**) Circular dichroism spectroscopy indicates similar structures of wild-type and I224 mutant Ddx49, plot with regions tabulated below. (**C**) Representative immunoblot of wild-type and I224 mutant Ddx49. (**D**) Representative helicase assay showing residual single stranded RNA in zebrafish I224N mutant zebrafish Ddx49, not present for wild-type protein. The human protein had less helicase activity than either zebrafish protein.

Since many DDX family proteins exhibit RNA helicase and ATPase activities (Gilman et al., 2017; Naineni et al., 2023; Dorner and Hondele, 2024; Tapescu and Cherry, 2024), we asked whether this was true of zebrafish Ddx49 and whether such activities were affected by I224N substitution. Wild-type Ddx49 exhibited robust helicase activity *in vitro*, whereas I224N had a marked reduction in activity compared to wild-type, albeit greater than observed for human wild-type DDX49 (**Figure 6C**; **Figure 6—figure supplement 1D**). Wild-type zebrafish Ddx49 also exhibited robust ATPase activity *in vitro*, though in this instance I224N mutant Ddx49 exhibited greater levels of ATPase activity than the wild-type; human wild-type DDX40 had greater ATPase activity than either zebrafish protein (**Figure 6D**).

### Ddx49 requirement for ribosome biogenesis

DDX49 of human and presumptive ortholog Dbp8 of *Saccharomyces* regulate pre-ribosomal RNA (pre-rRNA) processing and abundance, and Dbp8 is required for small ribosomal subunit assembly (Daugeron and Linder, 2001; Awasthi et al., 2018). We therefore sought to learn if zebrafish Ddx49 is similarly required for ribosome assembly by comparing 48-hour post-fertilization embryos that were either wild-type or injected with CRISPR/Cas9 reagent (AltR268; F0) targeting *ddx49*. Sucrose gradient profiling revealed peaks corresponding to small and large ribosomal subunits (40S, 60S), assembled ribosomes (80S) and polysomes in wild-type embryos; by contrast, *ddx49* F0 mosaics lacked discernible peaks for the small ribosomal subunit, had much-reduced peaks for mature ribosomes and polysomes, and a larger peak for the large ribosomal subunit (**Figure 7A,B**). These results imply a defect in small ribosomal subunit biogenesis, leading to a deficiency in assembled ribosomes and polysomes during translation, and surplus large subunit.

**Figure 7.**
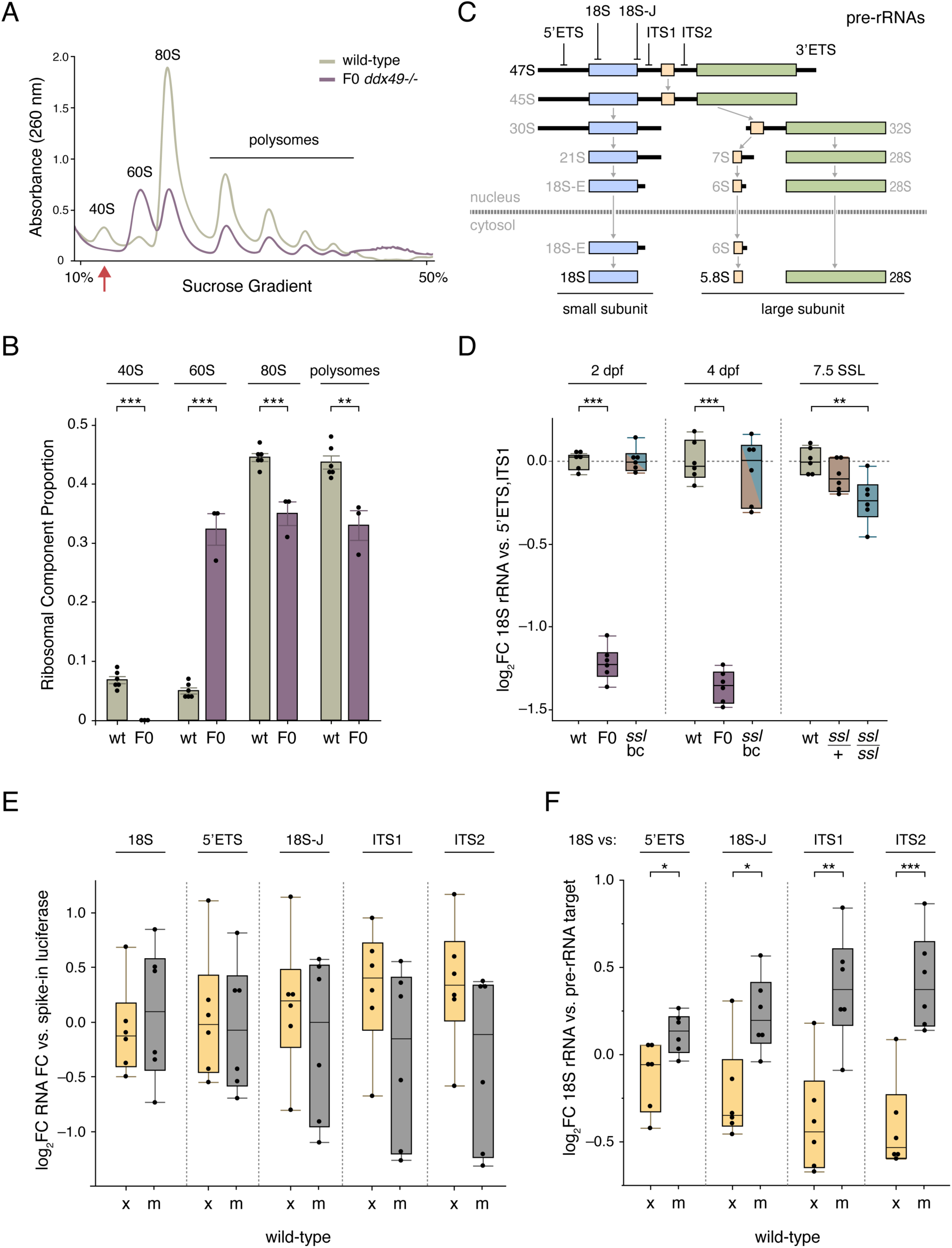
*ddx49* requirement for ribosome biogenesis. (**A**,**B**) Representative sucrose gradient profiles (**A**) and quantifications (**B**) for wild-type and *ddx49* CRISPR-targeted F0 embryos, with the latter exhibiting no 40S peak (arrow), reduced peaks for assembled ribosomes and polysomes and a superabundance of free 60S subunit. (**C**) Pre-rRNA processing for small and large ribosomal subunit rRNAs inferred for vertebrates, and simplified from (Henras et al., 2015). ⟘, qRT-PCR targets. (**D**) 18S rRNA target amplification relative to combined 5’ETS and ITS1 pre-rRNA targets; normalizations relative to pre-rRNA targets individually did not yield qualitatively different results. “*ssl* bc” represents pooled *ssl* backcross embryos (3:1 *ssl* to *+* alleles). (**E**) Relative quantities of rRNA targets in FAC-sorted xanthophores and melanophores of wild-type fish at 12 SSL. All cell-type differences *P*>0.1 by ANOVA. (**F**) Xanthophores had lower levels of 18S rRNA relative to pre-rRNA targets than did melanophores isolated from wild-type fish at 7.5 SSL. ***, *P*<0.0001; **, *P*<0.001, *, *P*<0.05 by ANOVA.

**Figure 7—figure supplement 1.**
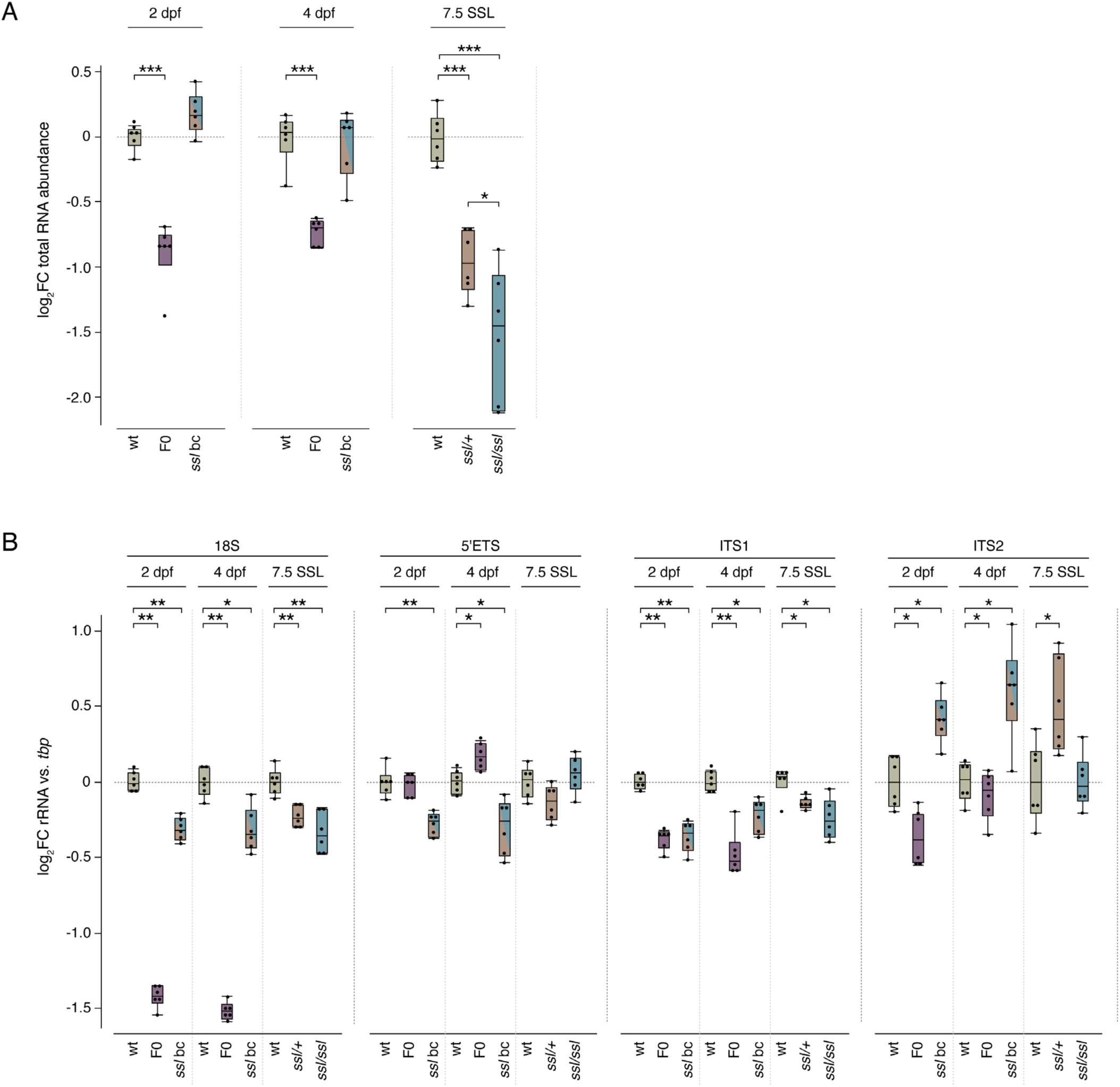
Reduced RNA abundance and defective rRNA processing due to *ddx49* mutation. (**A**) Total RNAs isolated from mutants for *ddx49* were less abundant than controls, both for mosaic CRISPR-targeted *ddx49* F0 embryos and for heterozygous and homozygous *ssl* mutants compared to wild-type fish of the same genetic background during later larval development. Controls were wild-type (wt) fish of the same genetic background as *ssl* (AB) and were, for comparisons at 2 days post-fertilization (dpf), the uninjected siblings of CRISPR-injected individuals. The substantial reduction in total RNA abundance for *ssl/+* compared to wt at 7.5 SSL is notable for a phenotypically recessive allele and could indicate *bona fide* differences in total RNA abundance per cell or developmental differences in cell size or number compared to homozgous wt controls (see Discussion). (**B**) Pre-rRNA targets varied significantly in abundance between controls and *ddx49* mutants at the same stage, both for targets predominantly representing “pre-18S” processing (5’-ETS, ITS1) and “pre-5.8S/28S” (ITS2; see Figure 7C) indicating an overall dysregulation of pre-rRNA processing. Notably, 18S target abundance was reduced in *ssl* homozygotes at 7.5 SSL and pooled *ssl* backcross embryos (“*ssl* bc” with 3:1 *ssl* to *+* alleles) at both 2 dpf and 4 dpf, in contrast to comparisons of 18S abundance to pre-rRNA targets shown in Figure 7D. We did not seek to resolve this difference given the clear indications, overall, of defects in pre-rRNA processing. Comparisons of means to wt controls of the same stages used Dunnet’s Method and a *post hoc* comparison total RNA between *ssl/+* and *ssl/ssl* at 7.5 SSL in A used Tukey-Kramer. *, *P*<0.05; **, *P*<0.01; ***, *P*<0.001.

**Figure 7—figure supplement 2.**
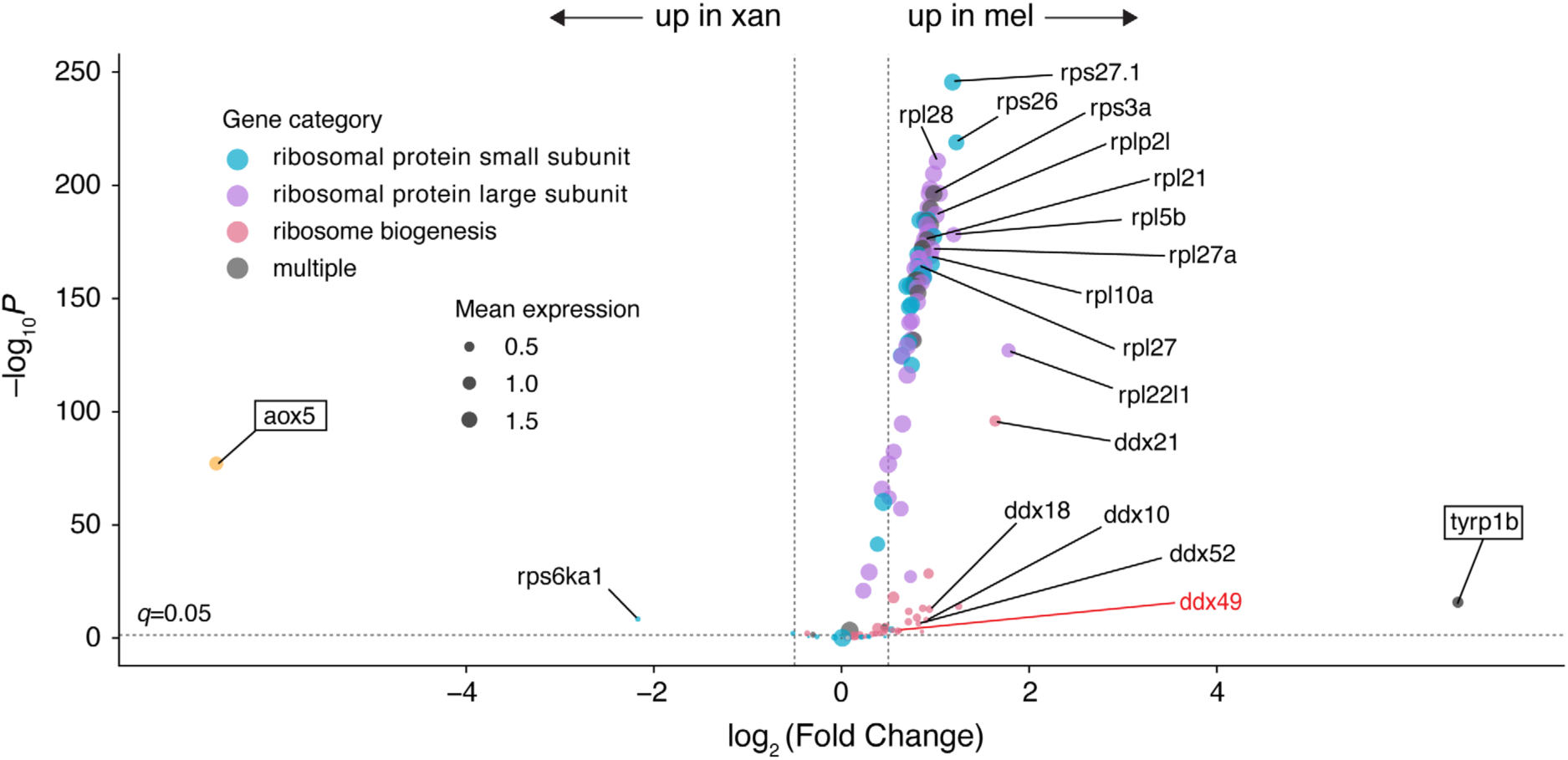
Differential expression of genes encoding cytosolic proteins of small and large ribosomal subunits and other genes required for ribosome biogenesis (**Supplementary file 1—Table 1**) between xanthophores and melanophores of wild-type fish. Xanthophore and melanophore specific markers *aox5* and *tyrp1b* are shown as benchmarks. (Full results in **Supplementary file 1— Table 7**.)

We predicted the *ssl* mutant allele would also lead to ribosomal subunit imbalance and ribosome deficiency, albeit of probably lesser magnitude than CRISPR-targeted F0 alleles. Yet preliminary gradient profiles for whole 7.5 SSL larvae did not reveal defects, nor did profiles for pooled embryos at 48 hpf, derived from backcrosses of male *ssl/ssl* to female *ssl/+* and so having 3:1 mutant to wild-type alleles. It was not feasible to PCR-genotype enough embryos to generate homozygous pools, nor was it possible to isolate enough xanthophores from older larvae to assay only the most affected cell type. Given roles for Ddx49 orthologs in rRNA processing, we turned to quantitative PCR to determine if *ssl* mutants have alterations in the abundance of pre-rRNA or rRNA transcripts likely to affect ribosome subunit biogenesis.

Fully processed 18S rRNA of the small ribosomal subunit and 5.8S and 28S rRNAs of the large ribosomal subunit derive from larger 47S pre-rRNA that undergoes a series of endonucleolytic cleavages and exonuclease digestions (**Figure 7C**). To learn if these steps are defective in *ddx49* mutants we designed qRT-PCR primers to target sites within the 5’ external transcribed spacer (5’ETS), internal transcribed spacers 1 and 2 (ITS1, ITS2), within the 18S rRNA itself, and spanning the junction between 18S and flanking precursor sequence (18S-J). Because the spacer sites are lost during pre-rRNA maturation we could evaluate processing efficiency of small subunit 18S rRNA by comparing its abundance to that of non-retained sites, and by comparing all rRNA amplicons to non-rRNA controls.

Isolation of RNAs for qRT-PCR comparisons revealed deficiencies in total RNA quantities as compared to wild-types of the same stages, both for CRISPR F0 and *ssl* heterozygotes and homozygotes (**Figure 7—figure supplement 1A**), consistent with defects in abundance of rRNAs, which typically constitute 80–85% of total RNA (Green and Sambrook, 2020). Aligning with the reduced small ribosomal subunit abundance above, CRISPR F0 embryos had significantly less 18S target amplification relative to combined 5’ETS and ITS1 targets, as compared to wild-type embryos (**Figure 7D**). And consistent with a defect in small subunit production owing to I224N substitution during later larval development, *ssl/ssl* mutants likewise had reduced ratios of 18S to 5’ETS/ITS1, in comparison with *ssl/+* siblings at the same stage; these defects were not apparent in comparisons of pooled *ssl/+* and *ssl/ssl* embryos to wild-type at 2 dpf or 4 dpf. We additionally compared rRNA targets to a non-rRNA control gene, *TATA-box binding protein* (*tbp*), after pre-screening candidate control genes not differentially expressed in *mitfa*:Eos+ cells by scRNA-Seq (log2FC=–0.07 in *ssl*/*ssl* compared to *ssl/+*; *P*=0.06), which was not the case for other commonly used control transcripts (e.g., *actb1*: log2FC=– 0.39, *P*=3.0E-33). Using *tbp* as control for normalization revealed significant reductions in 18S rRNA amplicon in *ddx49* mutants, with both decreases and increases in precursor rRNA targets relative to same-stage wild-types (**Figure 7—figure supplement 1B**). Together these findings indicated dysregulation of rRNA processing and defects in 18S rRNA maturation with *ddx49* mutations and the likelihood of diminished small ribosomal subunit biogenesis in *ssl* larvae, as in CRISPR-targeted *ddx49-/-*F0 mosaics.

Given these effects on ribosomal rRNA processing and ribosome biogenesis, we asked whether cell-type specific differences in these processes in wild-type fish might make xanthophores more vulnerable than melanophores to the *ssl* Ddx49 mutation and so isolated these cells by FACS using *aox5*:PALM-EGFP and *tyrp1b*:PALM-mCherry transgenes, respectively (McMenamin et al., 2014). By qRT-PCR normalized to a spike-in control, xanthophores did not differ significantly from melanophores in absolute abundances of 18S amplicon, representing mature and immature transcripts, nor did precursor rRNA targets differ clearly in abundance, including 18S-J, which overlapped the junction between 18S and precursor transcript (**Figure 7C,E**); a trend towards increased abundance of 3’ amplicons did not reach statistical significance (e.g., *P*=0.11 for ITS2). By contrast, ratios of 18S rRNA to each of the four pre-rRNA targets were significantly less in xanthophores than melanophores (**Figure 7F**), resembling the overall deficiency in *ssl/ssl* compared to *ssl/+* at the same stage (**Figure 7D**). To assess expression of genes encoding ribosomal proteins or biogenesis factors, we subsetted the scRNA-seq dataset for 124 such genes in xanthophores and melanophores of phenotypically wild-type *ssl/+* fish: 101 of these transcripts were more abundant in melanophores than in xanthophores (*q*<0.05, log2FC>0.5) (**Figure 7—figure supplement 2**).Together these observations suggest that a pre-existing state of rRNA processing or homeostasis in wild type xanthophores may be exacerbated past a threshold for phenotypic penetrance by Ddx49 dysfunction.

### *ssl* phenotypes are Tp53-dependent

Defects in ribosome biogenesis reduce translational activity and elicit nucleolar stress that is manifested in part through Tp53, leading to ribosomopathies that are tissue- or cell-type specific yet sometimes rescuable by loss of *tp53* (Danilova and Gazda, 2015). Central to this response is the E3 ubiquitin ligase Mdm2, which targets Tp53 for degradation in a manner facilitated by the non-canonical cyclin Ccng1 (Kimura and Nojima, 2002; Okamoto et al., 2002); ribosomal proteins such as Rps27l and Rpl11, when not incorporated into ribosomes, sequester Mdm2 and thereby stabilize Tp53, promoting cell cycle arrest (Xiong et al., 2014; Xu et al., 2023; Lindstrom, 2025). Distinct from this pathway, *cdkn2a/b* is the single teleost locus ancestral to mammalian *CDKN2A* and *CDKN2B*. The product of *cdkn2a/b* is INK4-like, blocking cell cycle progression through Cdk4/6 and promoting senescence, though unlike mammalian ARF it does not stabilize Tp53 (Sharpless and Sherr, 2015; Regneri et al., 2019). As noted above, all of these loci were upregulated in xanthophores of *ssl* mutants (**Figure 3C**), which also had reduced signatures of active translation and increased signatures of both nucleolar stress marker genes and Tp53 transcriptional targets. Mutational effects on the latter two signatures were more severe for xanthophores than melanophores, consistent with differential phenotypic severity at the cell-population level (**Figure 8A–C**).The fold-increase in transcript abundance owing to *ssl* homozygosity for a representative Tp53 target, *mdm2*, was markedly greater in xanthophores than melanophores (log2FC=2.7 and 1.0, respectively; difference in response magnitude between cell types: *P*=2.99E-12). Transcripts for *tp53* itself were significantly increased in xanthophores of *ssl* mutants compared to *ssl*/+ siblings (log2FC=0.94, *q*=1.1E-8) whereas the reverse was observed in melanophores (log2FC=–0.39, *q*=8.1E-3; difference in response between cell types, *P*=3.71E-13), suggesting that relatively modest increases in Tp53 target gene transcript abundances in melanophores reflected post-transcriptional rather than transcriptional regulation of *tp53*.

**Figure 8.**
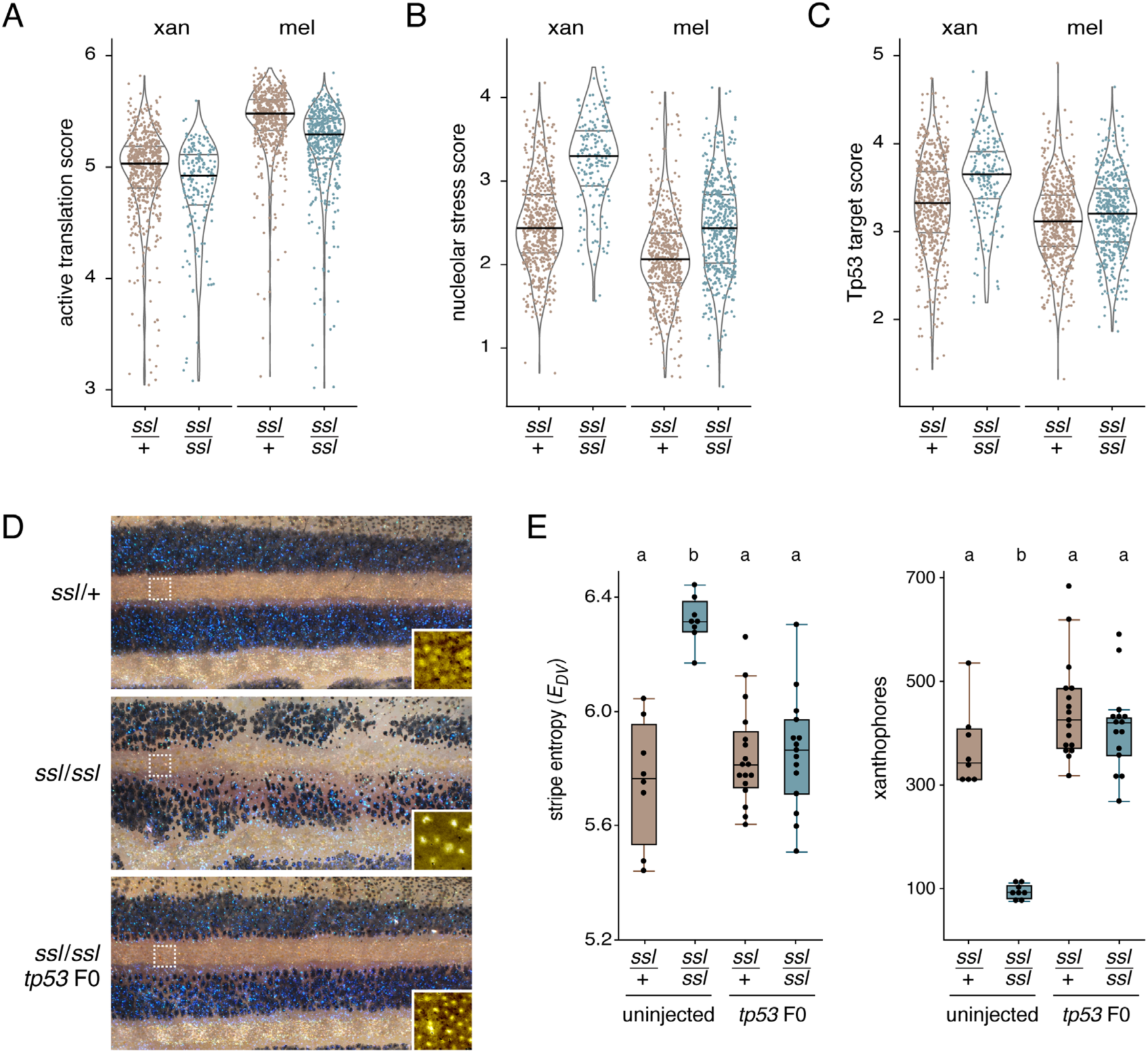
Tp53-dependence of *ddx49* mutant phenotypes. (**A**) Transcriptomic indicators of active translation (a subset of genes associated wirh translation overall; **Supplementary file 1—Table 1**) were reduced in both xanthophores and melanophores of *ssl/ssl* mutants compared to wild-type siblings. (**B**,**C**) Signature scores for marker genes of nucleolar stress as well as transcriptional targets of Tp53 activation, were both elevated in cells of ssl/ssl mutants, and at higher levels in xanthophores than in melanophores. Signature gene sets omitted loci encoding structural components of the ribosome itself and shared some loci that mark both nucleolar stress and represent direct targets of Tp53. (**D**,**E**) *ssl* mutant pigmentary defects were rescued in *ssl/ssl* mutants by mosaic CRISPR-targeting of *tp53*. Brightfield images illustrate melanophore pattern and insets show pseudocolored xanthophores at higher magnification. Stripe entropy (*EDV*) measures regularity of stripe pattern (McCluskey et al., 2021). Shared letters above plots indicate means not significantly different (*P*>0.05) in Tukey-Kramer *post hoc* comparison (overall for stripe entropy and xanthophore number, *F*3,44=16.9, 36.6; both, *P*<0.0001). Melanophore numbers at the stage examined for phenotypic rescue did not differ significantly across heterozygotes and homozygotes, regardless of *tp53* status (overall *F*3,44=2.5, *P*=0.07).

**Figure 8—figure supplement 1.**
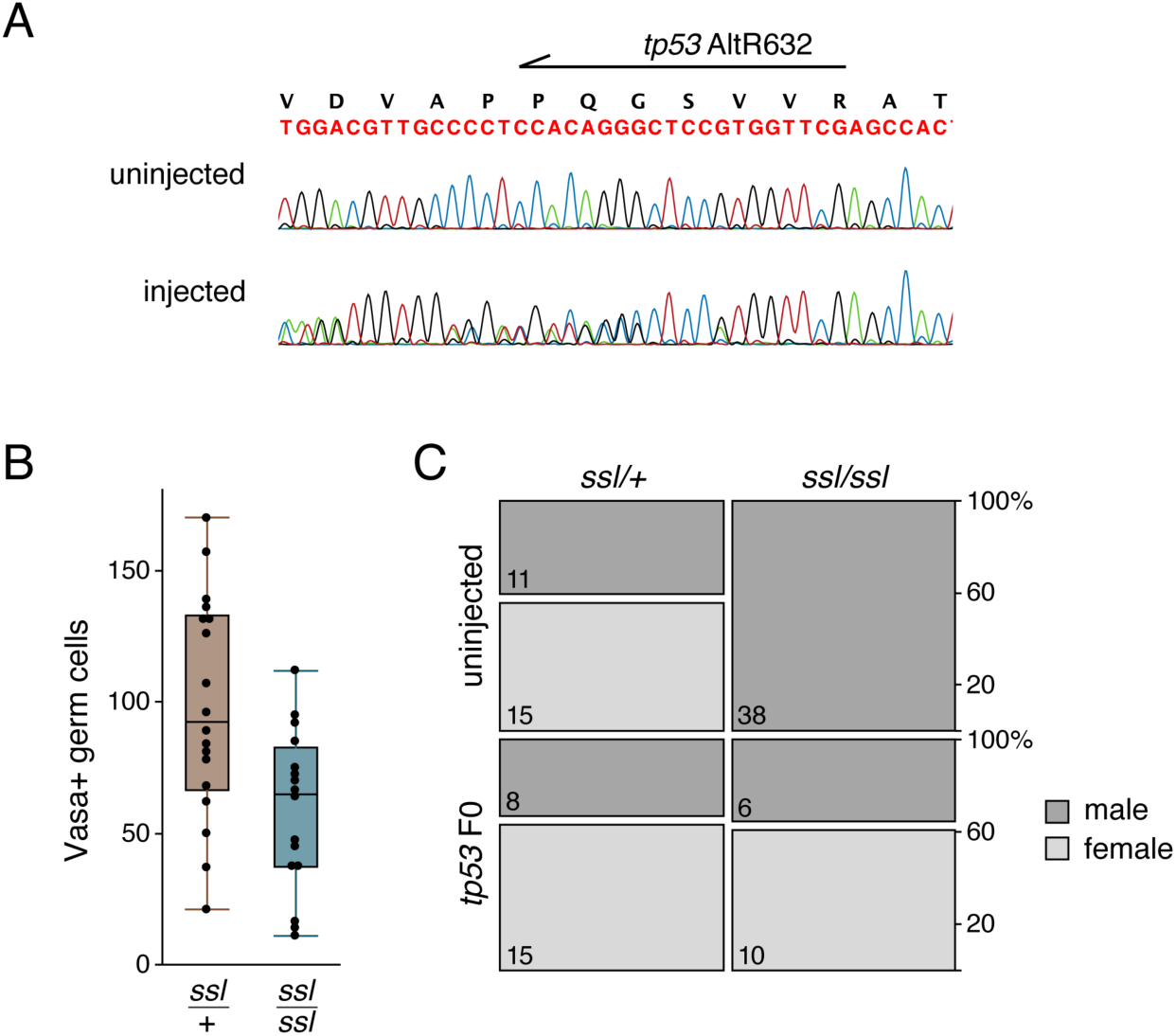
Tp53-dependence of *ssl* mutant phenotypes. (**A**) Representative sequencing across the *tp53* target site illustrating mutagenesis in a genetically mosaic adult fish. (**B**) Male-bias of *ssl/ssl* mutant sex ratio correlated with fewer germ cells compared to *ssl/+* type siblings (Wilcoxon S=203.5, *P*=0.009; *n*=34 larvae at 7–8 SSL). (**C**) Mosaicism for *tp53* mutations was sufficient to rescue female sex differentiation, restoring in *ssl/ssl* the same sex ratio observed for *ssl/+* siblings. The plot illustrates the 2 x 2 design, split vertically to show whether fish were uninjected controls—wild-type for *tp53*—or F0 mosaics for *tp53* mutations, and split horizontally show whether fish were heterozygous or homozygous for *ssl*. Dark grey represents percentage of males whereas light grey represents percentage of females within each quadrant; numbers within boxes are counts of adult fish. Uninjected fish that were *ssl/ssl* developed only as males (upper right), whereas fish mosaic for *tp53* mutations that were ssl/ssl developed as both males and females (lower right; *tp53* condition x *ddx49* genotype interaction, χ^2^=17.2, d.f.=1, *P*<0.0001).

To learn if *ssl* mutant phenotypes are Tp53-dependent we targeted the locus by CRISPR/Cas9 injection. Pigmentary phenotypes of *ssl/ssl* individuals mosaic for *tp53* mutations were indistinguishable from wild-type, a rescue evident in both recovery of stripe pattern and xanthophore numbers (**Figure 8D,E**; **Figure 8 — figure supplement 1A**).

The survival of *ssl/ssl* mutants through reproductive maturity, and their nearly exclusive development as males also afforded an opportunity to assess *tp53*-dependence of sex differentiation in this background. Male zebrafish develop fewer primordial germ cells than females and genetic or experimental depletion of these cells results in female-to-male sex reversal (Rodriguez-Mari et al., 2010; Tzung et al., 2015). In *ssl/ssl*, Vasa+ germ cells were significantly fewer than in *ssl/+* siblings (**Figure 8 — figure supplement 1B**). Indicating a Tp53-dependence of biased sexual differentiation, *ssl/ssl* fish wild-type for *tp53* developed only as males, whereas *ssl/ssl* fish mosaic for *tp53* mutations were as likely as their *ssl/+* siblings to develop as females (**Figure 8 — figure supplement 1C**).

## Discussion

We have shown that a new pigment pattern mutant of zebrafish, *sisley* (*ssl*), results from a viable missense allele of *ddx49*, encoding a DEAD-box RNA helicase studied previously only in yeast and cell culture. Zebrafish Ddx49 is required for ribosome biogenesis, and the *ssl* allele leads to proliferative arrest of a neural crest derived pigment cell, the yellow/orange xanthophore, with far milder effects on a lineally related cell, the melanophore. Defects in xanthophore abundance and pigment pattern, and a bias in sexual differentiation involving an independent lineage, primordial germ cells, were both rescued by mutation of the tumor suppressor gene *tp53*. These findings provide insights into the cell-type specificity of ribosomopathies, roles for Ddx49 in vertebrate ribosome biogenesis, and how population expansion of particular cell types allows pattern formation in an accessible model of organogenesis.

Ribosomopathies have long been recognized for pathologies that are surprisingly cell- and tissue-type specific, given the ubiquity of ribosomes themselves (Yelick and Trainor, 2015; Mills and Green, 2017; Farley-Barnes et al., 2019; Kampen et al., 2020). This discordance prompted reasonable suggestions of extra-ribosomal functions for some ribosome-associated genes, and may have slowed recognition of ribosome dysfunction in pathologies where causal loci were not already known to have ribosomal roles (Group et al., 1996; Draptchinskaia et al., 1999; Ridanpaa et al., 2001; Boocock et al., 2003; Dixon et al., 2006; Yelick and Trainor, 2015). Yet even as more pathologies have been linked to ribosome dysfunction, no single mechanism has explained their specificities, which have instead been attributed to especially high developmental demands for translation (Falcon et al., 2022), cell-type differences in ribosome homeostasis and recycling (Mills et al., 2016; Mills and Green, 2017), deficient translation of lineage-critical mRNAs (Ludwig et al., 2014; Yang et al., 2016; Khajuria et al., 2018; Tiu et al., 2021), cell-type specific stress responses (Calo et al., 2018; Oyarbide et al., 2025), tissue-specific physiology or environment (Schneider et al., 2016; Yang et al., 2016; Joyce et al., 2019), lineage specific ribosomal modifications (Farley-Barnes et al., 2019; Genuth et al., 2022; Ni and Buszczak, 2023; Poverennaya et al., 2026), and in some cases combinations of these.

Our findings suggest how such specificity can arise even between two closely related cell types. *ssl* mutants had reduced total RNA and reduced 18S rRNA relative to 47S precursor rRNA, pointing to defects in precursor rRNA processing. Remarkably, xanthophores, far more affected by the *ssl* mutation than melanophores, also had considerably lower ratios of 18S rRNA to precursor rRNA even in the wild-type, as well as lower levels of transcripts for ribosomal proteins and ribosome biogenesis factors. This correspondence suggests that an inherent difference in rRNA processing and ribosome homeostasis underlies the differential vulnerabilities of these cells, with further impairment of 47S processing and 18S maturation in *ssl* mutants pushing xanthophores into an overtly pathological state, evident as nucleolar stress, induction of Tp53 target genes and Tp53-dependent proliferative arrest.

We speculate that this difference reflects, at least in part, the differing developmental histories of these cells. This and a previous study indicate that most xanthophores of the young adult derive from early larval xanthophores that differentiate in the embryo, then lose their pigment and proliferate in a cryptic state before redifferentiating in a thyroid hormone-dependent manner (Le Guyader and Jesuthasan, 2002; McMenamin et al., 2014; Saunders et al., 2019). In some systems, ribosomal gene expression and biogenesis are curtailed as cells differentiate (Woolnough et al., 2016; Chau et al., 2018; Le Goff et al., 2021), and our findings lead us to suspect the same is true of xanthophores. If thyroid hormone “reawakens” rRNA transcription and ribosome biogenesis, as it can elsewhere (Tata, 1967), it does not seem to drive these processes in xanthophores to the levels of melanophores. By contrast, most adult melanophores differentiate from latent, multipotent progenitors during the larva-to-adult transition, with extensive migration and proliferation, and changes not only in melanization but also in cell size and shape, over just a few days (Parichy and Turner, 2003b; Budi et al., 2011; Dooley et al., 2013; Singh et al., 2016). Xanthophores may thus lack sufficient buffering of ribosome numbers or biogenesis to withstand Ddx49 dysfunction, whereas melanophores, more recently derived from progenitors, may be better equipped; indeed, variable expressivity of the *ssl* mutant phenotype could reflect differential regulation of the pattern, dependent upon the degree of population expansion achieved by the progenitor-derived pool of xanthophores. Testing these ideas should provide insights into how differentiation trajectories, and thyroid hormone, influence ribosome production and translational capacity.

Embryos mosaic for *ddx49* loss-of-function mutations had gross reductions in 18S rRNA, reduced ratios of 18S to precursor rRNA, and loss of the small ribosomal subunit. These findings agree with roles for DDX49 inferred from human cell lines in precursor rRNA processing (Awasthi et al., 2018) and ribosomal small subunit assembly (Tse et al., 2026). They also agree with studies of yeast, in which the homologue Dbp8 promotes cleavage of precursor rRNA at sites flanking 18S, and its depletion eliminates the small subunit and leaves a surplus of large subunit (Daugeron and Linder, 2001; Granneman et al., 2006), as we observed. Given that rRNAs constitute 80–85% of total RNA, reduced total RNA in mutants is consistent with DDX49 promoting rRNA transcription directly (Awasthi et al., 2018), or indirectly through translational insufficiency, or fewer cells owing to reduced rates of proliferation, cell cycle arrest or apoptosis.

Beyond reducing protein abundance, the *ssl* allele separates Ddx49 helicase and ATPase activities. The I224N substitution replaces a conserved hydrophobic residue at the N-terminal edge of the RecA2 domain, six amino acids beyond the interdomain linker between RecA1 and RecA2. We speculate that this substitution interferes with the ability of these domains to close upon or stably engage RNA, consistent with the sharply reduced helicase activity of the mutant protein *in vitro*. During productive cycles of RNA remodeling by DEAD-box proteins, binding of ATP is believed to promote closure of RecA1 and RecA2 around double-stranded RNA and conformational changes that separate the strands; ATP hydrolysis then releases the helicase from RNA for new cycles of remodeling (Jankowsky, 2011; Bohnsack et al., 2023; Dorner and Hondele, 2024). The increased ATPase activity of *ssl* Ddx49 thus implies a decoupling of ATP cycling from RNA remodeling. Similar decoupling has been observed for translation initiation factor eIF4A1 carrying mutations in the SAT motif, which abolished helicase activity while enhancing ATPase activity (Pause and Sonenberg, 1992). Strikingly, hydrophobic to non-hydrophobic substitutions within the interdomain linker itself led to similar outcomes, with the greatest losses in helicase activity accompanied by the greatest gains in ATPase activity (Lu et al., 2015). Most other mutations to DEAD-box proteins reduce or abolish both activities together (Sengoku et al., 2006; Elles and Uhlenbeck, 2008; Banroques et al., 2010) or, rarely, increase both (Rocak et al., 2005). The decoupled activities of *ssl* Ddx49 may thus provide a useful *in vivo*, adult-viable model for understanding DEAD-box protein function across vertebrate cell types. A recent study of human DDX49 points to an additional nuclease activity and RNA-binding interactions associated with the C-terminal regions and it would be interesting to assess such activities *in vivo* with zebrafish (Parkes et al., 2024). In that study and others, it has not been possible to obtain cell lines or yeast strains entirely lacking *DDX49* or its homolog (Daugeron and Linder, 2001) indicating essential function for cell viability, and the locus has not been associated with human pathologies (https://omim.org/entry/621419), consistent with early lethality of major-effect mutations.

Xanthophore deficiency and male-biased sexual differentiation were both rescued by *tp53* mutation, as are some ribosomopathy phenotypes in other systems (Barlow et al., 2010; Danilova and Gazda, 2015; Mills and Green, 2017; Calo et al., 2018; Watt et al., 2018; Tiu et al., 2021; Falcon et al., 2022). In xanthophores, transcriptomic changes pointed to a nucleolar stress response (Liu et al., 2016; Lafita-Navarro and Conacci-Sorrell, 2023), presumably underlying the proliferative arrest, without cell death, that we observed by fate mapping. How germ cells are depleted in *ssl* mutants remains unclear.

Deficiencies of primordial germ cells cause female-to-male sex reversal of zebrafish in other contexts, and in mutants for the Fanconi anemia gene *fancl*, simultaneous loss of *tp53* rescues these cells from apoptosis (Rodriguez-Mari et al., 2010; Tzung et al., 2015; Orban et al., 2021). Whatever the proximate cause of germ cell deficiency in *ssl*, it is notable that xanthophores and primordial germ cells—despite their very different states of commitment—share large size and the demands of rapid proliferative expansion and considerable biosynthetic output during a defined window (Percharde et al., 2017; Razmi et al., 2026). Indeed, germ cell numbers are much reduced, with hyperactivation of *tp53*, upon depletion of the pre-rRNA processing factor Puf-A in both zebrafish and mouse (Qiu et al., 2014; Ko et al., 2022). It will be interesting to learn whether *ssl* germ cells fail to proliferate, fail to survive, or both, and whether cell-type specific defects arise in other tissues, given the broad expression of *ddx49* and the slow growth and development of these fish.

Finally, our study underscores the utility of pigment pattern as an emergent read-out of lineage-specific population expansion, and the ability of developmental systems to compensate, or not, for deficits in essential cellular activities. Pigmentation has long been used to study mechanisms of pattern formation, but it also provides an especially accessible indicator of cell behaviors relevant far beyond stripes and spots. Indeed, mutations in ribosomal protein genes of mouse can increase or decrease pigmentation through effects autonomous or non-autonomous to the melanocyte lineage (Oliver et al., 2004; McGowan et al., 2008; Terzian et al., 2011; Watkins-Chow et al., 2013). Here, cell-autonomous dysfunction of a “housekeeping” gene yielded a phenotype resembling *csf1ra* mutants, with variable expressivity suggesting that *ssl* acts near a threshold for phenotypic penetrance and recalling the variable penetrance and expressivity of ribosomopathies in humans and their background dependence in animal models (Dixon and Dixon, 2004; Carlston et al., 2017; Fitriasari et al., 2025). In the contexts of both pattern formation and essential cellular mechanisms, we anticipate that forward genetic screens for pigmentary phenotypes will continue to provide useful and novel insights.

## Materials and methods

### Key resources table

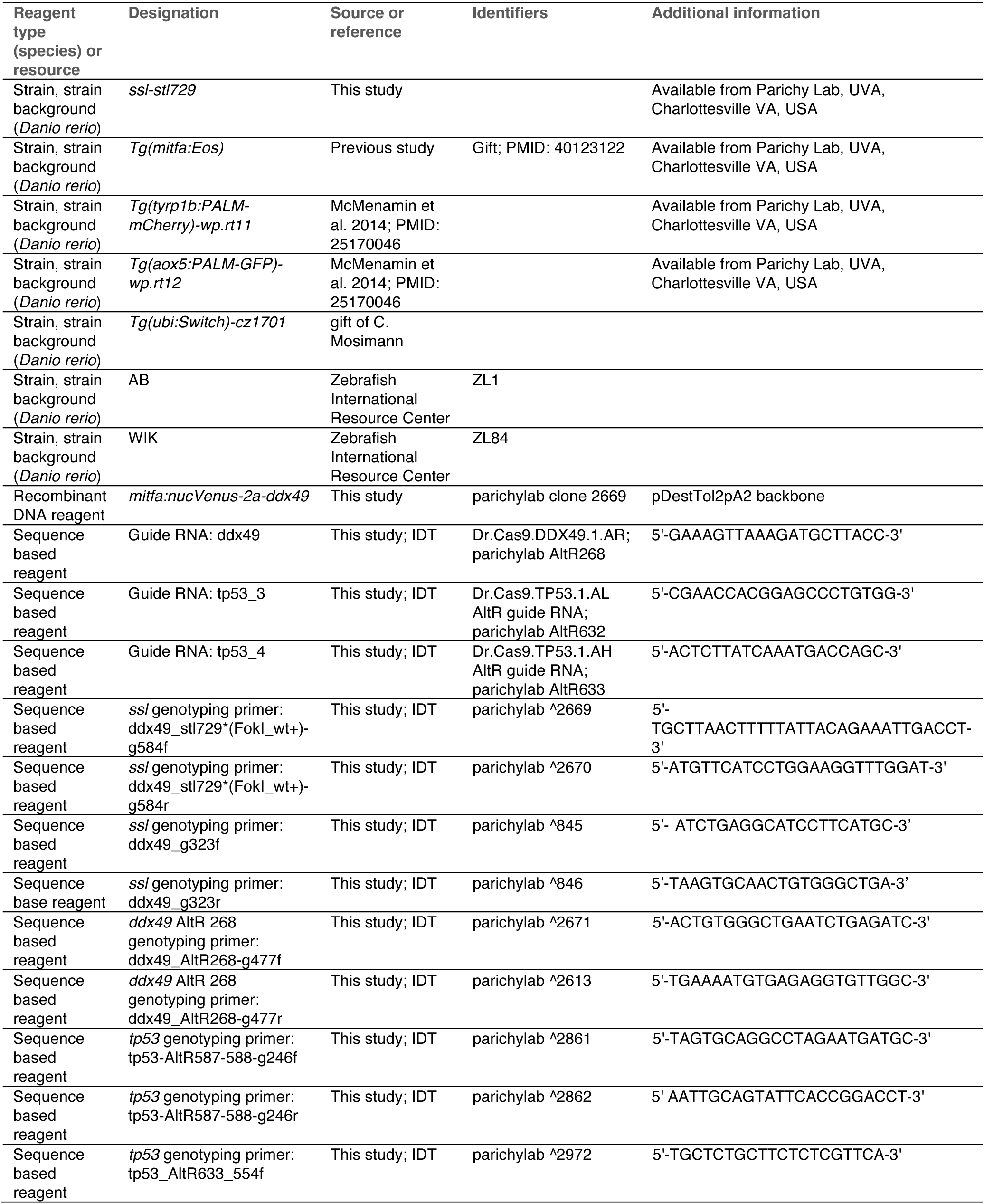
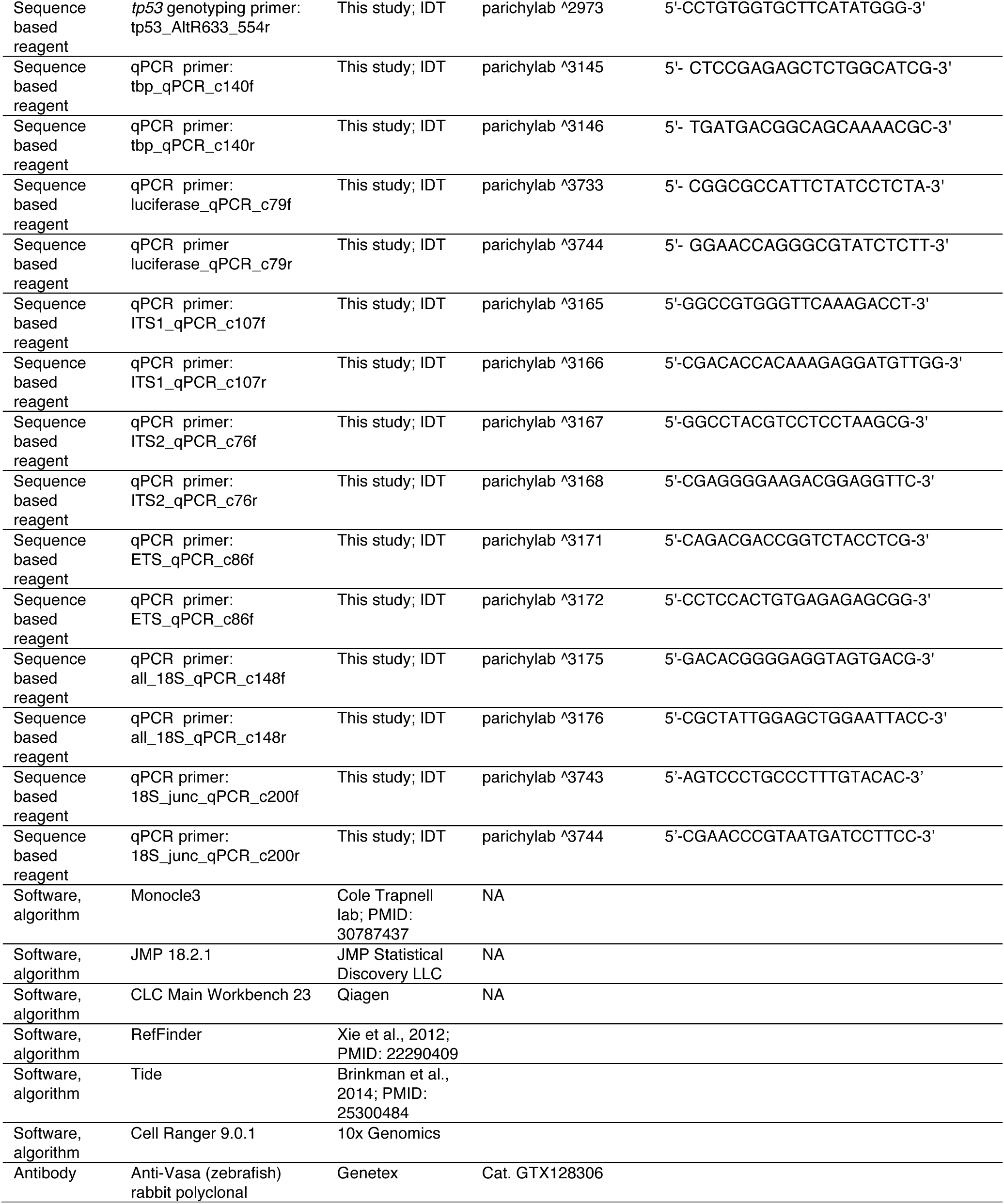

### Zebrafish husbandry, stocks and genotyping

Fish were reared at ∼28°C (14L:10D) with post-embryonic staging using standardized standard length in mm (SSL) (Parichy et al., 2009). Larvae were reared on marine rotifers enriched with Algamac (Reed Mariculture), *Artemia* and flake food; adults were fed *Artemia* and flake food. The *stl729* N-ethyl-N-nitrosourea allele was isolated in a forward genetic screen in the AB genetic background by standard methods (Solnica-Krezel et al., 1994) and was thereafter maintained in this background. *Tg(mitfa:Eos)* was generously provided by James Lister (ret.)(Korzeniwsky et al., 2025). To distinguish between wild-type and *ssl* mutant alleles we used PCR with primers 2669, 2670 ddx49_stl729*(FokI_wt+)-g584f,r (see Key resources table) and genotyping by FokI digestion, yielding fragments of 501, 45, 38 bp for wild-type and 546, 38 bp for *ssl* alleles, or primers 845, 846 ddx49-g323f,r for genotyping by Sanget sequencing. Comparisons of xanthophores and melanophores from wild-type used FAC-sorted cells from doubly transgenic fish for recombineered BAC transgenes *Tg(aox5:PALM-GFP)^wp.rt12^* and *Tg(tyrp1b:PALM-mCherry)^wp.rt11^* (McMenamin et al., 2014).

Animals were handled according to procedures approved in Institutional Animal Care and Use Protocol #4170 of the University of Virginia. Euthanasia was performed with overdose of MS222 (Syndel) followed by physical maceration.

### Imaging

Ontogenetic pigment phenotypes of *ssl* mutants and *ssl/+* siblings were assessed in three ways. Except where otherwise noted, melanosome and carotenoid vesicles were contracted toward cell centers by brief treatment with epinephrine, and fish were anesthetized with tricaine prior to imaging. Genotypes were confirmed by PCR followed by FokI digest and/or Sanger sequencing (see above). First, individual fish were imaged daily from 6.7–12.0 SSL as previously described (Patterson and Parichy, 2013), on a Zeiss AxioObserver microscope equipped with Zeiss Axiocam cameras and ZEN Blue software.

Melanophores and xanthophores were counted in a region ventral to the horizontal myoseptum, extending from the anterior margin of the dorsal fin to the posterior margin of the anal fin (Fig. 1C). Pseudocoloring of xanthophores for easier visualization followed (Korzeniwsky et al., 2025). Second, to assess developmental progression of pigment phenotypes and precursors without repeated handling, fish ∼6.0-12.0 SSL were randomly sampled from *ssl/+* x *ssl/ssl* crosses, expressing *Tg*(*mitfa:Eos)* and imaged once using both brightfield and epifluorescence illumination on a Zeiss AxioObserver. Eos+ xanthophores, melanophores and undifferentiated precursors were counted in the same region used in repeated imaging. To assess variability in adult patterns, juvenile fish were imaged with and without epinephrine treatment at 15.0 SSL on a Zeiss AxioZoom microscope. Tp53 F0s and uninjected control were imaged with and without epinephrine treatment on a Zeiss AxioZoom microscope with Axiocam color camera.

### Photoconversions

Photoconversions of *ssl* mutants and *ssl/+* siblings transgenic for *mitfa:Eos* were performed using a Zeiss LSM 800 scanning laser confocal with a 405 nm laser and ZEN blue software. Prior to Eos photoconversion and imaging, fish were anesthetized using tricaine. To examine proliferation of cryptic xanthophores, 1–2 cells were targeted for photoconversion per fish (6.1-7.0 SSL). Target cells were identified based upon morphology and location within the presumptive primary interstripe, in the region bordered by the dorsal and anal fins. To assess the contribution of latent precursor derived xanthophores, a region of the flank, posterior to the anus and extending from the dorsal to ventral margins of the body, was photoconverted at 6.3 – 7.1 SSL. Following photoconversion, fish were allowed to recover for 1-3 hours, then treated with epinephrine and imaged using a Zeiss Axio Observer equipped with a Yokogawa CSU-X1M5000 laser spinning disc and appropriate cameras to confirm that photoconversion was complete and restricted to target cells or region. Fish were then reared individually on flowing water and imaged 7 days after photoconversion.

### Single cell RNA-Sequencing

For scRNA-Seq skins were removed from *ssl/+* and *ssl/ssl* siblings at 12 SSL, pooled by genotype (*n*=15 per pool) and dissociated in Liberase TM (Roche) at 28 °C for 20 min, then filtered through a 70µm strainer. Healthy cells were sorted by fluorescence activated cell sorting for mitfa:Eos expression using a Cytek Aurora CS: debris were removed from the FSC/SSC plot, and singlets were further isolated using both FSC-H/FSC-A and SSC-H/SSC-A gates, followed by live/dead cell gating based on DAPI staining. Finally, Eos-positive cells were sorted based on the B1-A and B2-A channels. Libraries were produced with a 10X Genomics Chromium controller and Chromium Next GEM Single Cell 3’ Kit v3.1 with quality control and quantification assays using a Qubit fluorometer (Thermo) and a 4150 TapeStation (Agilent). Libraries were sequenced with NovaSeq X Plus at Novogene then processed for bar codes and unique molecular identifiers (UMI) using Cell Ranger 9.0.1 and mapped using zebrafish genome annotations filtered for protein-coding genes (Lawson et al., 2020) and presumptive doublets removed. Data were analyzed with Monocle3 (Cao et al., 2017) following (Saunders et al., 2019; Huang et al., 2025) after filtering for mitochondrial reads (<2%), total UMI (>500) and number of genes expressed (>250). After dimensionality reduction by Uniform Manifold Approximation and Projection (UMAP) and Louvain clustering cell groups were assigned to types using marker transcripts previously defined (Saunders et al., 2019; Aman et al., 2023). Differential expression analyses were performed either on subsets of the full dataset or using linear contrasts defined to isolate main effects of genotype and cell-type specific effects independent of cell-type representation, using a model of gene expression=∼genotype * cell-type.

### Mapping the *ssl* mutation

Bulk segregant analysis was performed similar to the method outlined in (Bowen et al., 2012) with modifications to account for not having established marker SNPs from sequenced parental strains. Large, full sibling backcrosses were sorted for the *ssl* phenotype and fin clips were taken for genomic DNA isolation from the two pools. Reads were sequenced on an Illumina NextSeq550 and aligned to the zebrafish genome with bwa mem from BWA v0.7.17 (Li and Durbin, 2009) using default parameters. SNPs were called using HaplotypeCaller in GATK v4.0.0.0 (Poplin et al., 2018) with the following parameters: ’--sample-ploidy 2 --dont-use-soft-clipped-bases true --padding-around-indels 5 --padding-around-snps 0 --native-pair-hmm-threads 2 --heterozygosity 0.01’. To identify high-quality mapping SNPs, sites with minor allele frequencies below 0.15 in the WT pool were removed. High quality mapping SNPs were scored for minor allele frequency and homogeneity (Bowen et al., 2012), which were then plotted in 100 SNP sliding windows across the genome. SNPs with high homogeneity within protein-coding genes were prioritized based on predicted effects from Ensembl’s Variant Effect Predictor (McLaren et al., 2016).

### RNA isolation and First-strand cDNA synthesis

To compare rRNA expression among genotypes and developmental stages, we collected samples from three stages, 2 dpf, 4 dpf, and 7.5 SSL, with six biological replicates per stage for each genotype assessed. For homogenization, 2 dpf and 4 dpf embryos were rinsed with 1× PBS and 25 individuals were transferred into 1 ml ice-cold TRIzol (Invitrogen) as one biological replicate for RNA isolation, then homogenized using a 23G needle and a 25G needle. For 7.5 SSL, fish were euthanized, heads removed and trunks rinsed with 1× PBS with four trunks per replicate transferred into 1 ml ice-cold TRIzol (Invitrogen) in a 2 ml Eppendorf LoBind tube with one 1/4” ceramic sphere (MP Biomedicals) and 50–100 ul of Lysing Matrix A (MP Biomedicals), then homogenized using a Tissuelyser II (Qiagen) at maximum speed for 30 s. Homogenized mixtures were either processed immediately for RNA isolation or stored at -80°C for long-term storage. Subsequent total RNA isolation and on-column DNase treatment were performed using Direct-zol RNA Miniprep Kits (Zymo Research) per manufacturer instructions. RNA concentration and purity were assessed using a DS-11+ spectrophotometer (DeNovix). First-strand cDNA was synthesized from 500 ng total RNA using a GoScript Reverse Transcriptase Kit (Promega) with priming by random hexamers; incubations at 50°C for 10 min and 55°C for 10 min were added after the recommended extension step at 42°C for 60 min. cDNA was diluted ten-fold as qPCR template.

For comparison of rRNA expression between xanthophores and melanophores, skins from 20 individual 12 SSL *Tg(tyrp1b:PALM-mCherry)*; *Tg(aox5:PALM-EGFP)* fish were removed and dissociated as described above per replicate, with six biological replicates total. Xanthophores and melanophores were sorted by fluorescence-activated cell sorting based on aox5:PALM-EGFP and tyrp1b:PALM-mCherry fluorescence, respectively, with a target of 2.5 × 10⁵ cells for each population. After sorting, 1.042 × 10² copies of Luciferase Control RNA (Promega) per sorted cell were added to each sample before lysis as an external spike-in reference for relative expression level calculations. Cell pellets were resuspended in 100 µl BL + 1-Thioglycerol (TG) Buffer and lysed by pipetting. Cell lysates were snap-frozen in liquid nitrogen and stored at −80°C until RNA purification. Total RNA was purified from cell lysates using the ReliaPrep RNA Miniprep System (Promega) according to the manufacturer’s protocol. At the end of the protocol, RNA was eluted in 25 µl nuclease-free water. A 16.5 µl aliquot of the purified RNA was immediately used for first-strand cDNA synthesis, while the remaining RNA was used for RNA quality and concentration assessment. All purified RNA samples had an RINe (RNA Integrity Number equivalent) above 8.8, as measured using the High Sensitivity RNA ScreenTape Assay (Agilent). Prior to first-strand cDNA synthesis, 1.186 × 10² copies of Kanamycin Positive Control RNA (Promega) per sorted cell were added to each sample as a second external spike-in reference. Reverse transcription was performed in a 40 µl reaction using GoScript Reverse Transcription Mix with Random Primers (Promega) under the conditions described above. Following reverse transcription, cDNA was diluted two-fold and used as the template for qPCR.

### Quantitative PCR

Prior to qPCR, Sanger sequencing confirmed that qPCR primer binding sites did not overlap with known or previously undetected sequence variants and primer efficiencies were determined using 1:2 serial dilutions of pooled cDNA from all samples. For each gene or rRNA target, two primer pairs were evaluated when possible, and the best-performing pair was selected for use. rRNA targets were compared internally, 18S (precursor and processed rRNA) relative to 5’ETS and ITS1 (precursor, using geometric mean expression of combined targets). Additionally, candidate control protein-coding genes (*actb1*, *gapdh*, *tbp*, *b2m*, and *ef1a*) were evaluated for expression stability using RefFinder (Xie et al., 2023) and by comparing genotype effects in scRNA-Seq with *tbp* was selected for normalization based on amplification efficiency and comparative insensitivity to *ddx49* mutation.

qPCRs used 4 µl diluted first-strand cDNA, 10 pmol each forward and reverse primer, and GoTaq qPCR Master Mix (Promega), with nuclease-free water added to a final reaction volume of 20 µl in a 96-well plate. Amplification was carried out on a 7500 Fast Real-Time PCR System (Applied Biosystems) with initial denaturation at 95°C for 10 min, followed by 40 cycles of 95°C for 20 s and 60°C for 60 s. After amplification, melting curve analyses verified PCR product specificities and that all samples had single peaks. Randomly selected samples were subjected to agarose gel electrophoresis to further confirm specificity. For 18S-J the two better-performing pairs showed two peaks in the melting curves, both above 80 °C. Because the amplicon is approximately 200 bp and the available primer binding sites are limited, we performed agarose gel electrophoresis for all samples to verify product specificity. All reactions were performed in triplicate, and only reactions with Ct standard deviations <0.2 among technical replicates were included in subsequent analyses. Ct values were defined as the cycle number at which fluorescence crossed the threshold above background defined by the instrument. Relative expression levels were normalized to luciferase or tbp and calculated using the Pfaffl method (Pfaffl, 2001), incorporating the experimentally determined amplification efficiencies of both target and reference genes.CRISPR/Cas9 mutagenesis

### CRISPR/Cas9 mutagenesis

Targeted knockout of *ddx49* and *tp53* in F0 mosaics used IDT AltR reagents with separate tracrRNA and guide RNAs injected into 1-cell stage embryos (∼1 nl 5 µM premixed RNAs:Cas9 complex) (Hoshijima et al., 2019) with mutagenesis efficiencies validated by Sanger sequencing and TIDE software (tide.nki.nl) (Brinkman et al., 2014). A single ddx49 guide RNA was used whereas two different *tp53* guide RNAs were used with similar efficiencies of mutagenesis (Key Resources table).

### Cell transplantations, rescue injections and cloning

Blastula stage transplants were performed as described (Eom et al., 2021). Cells from wild-type donors of transgenic line *Tg(ubi:Switch)^cz1701^* (Lalonde et al., 2022) were transplanted to *ssl/ssl* mutant hosts. Host embryos with EGFP+ cells were reared to juvenile stages and imaged using a Zeiss AxioZoom with monochrome and color Axiocam cameras. Constructs expressing wild-type *ddx49* cDNA under the control of a ∼2.2 kb fragment of the *mitfa* promoter (Eom et al., 2021) were generated by isothermal assembly using the pDestTol2pA2 vector (Kwan et al., 2007). One-cell embryos from crosses of *ssl/+* x *ssl/ssl* were injected with Tol2 mRNA and *mitfa:nucVenus-2a-ddx49* vector following standard procedures (Suster et al., 2009). Embryos with nucVenus+ cells at 24 hpf were reared to young adult stages and imaged on Zeiss AxioObserver or Zeiss AxioZoom microscopes using brightfield and epifluourescence illumination. To visualize nuclear Venus expression in pigment cells, fish were treated with epinephrine to contract pigment vesicles prior to imaging. Genotypes of injected fish were determined Sanger sequencing of PCR products (see above).

### Polysome gradient

Wild-type (WT) and *ddx49* F0 knockout embryos at 2 dpf were dechorionated with 1mg/ml Pronase (Sigma) in 10% Hank’s solution, and deyolked (Link et al., 2006; Shah et al., 2024): yolk sacs of dechorionated embryos were disrupted by pipetting with a P1000 pipettor in deyolking buffer (55 mM NaCl, 1.8 mM KCl, 1.25 mM NaHCO₃, 0.1 μg/mL cycloheximide, and 1× Roche cOmplete EDTA-free protease inhibitor cocktail). Samples were then shaken at 1100 rpm for 5 min to dissolve the yolk.

Embryos were pelleted at 300 × g for 30 s and washed twice with embryo wash buffer (110 mM NaCl, 3.5 mM KCl, 2.7 mM CaCl₂, 10 mM Tris/Cl, pH 8.5), with shaking at 1100 rpm for 2 min during each wash. A total of 650 or 750 embryos per genotype were lysed in 550 μl of lysis buffer (20 mM HEPES-KOH (pH 7.4), 100 mM KCl, 5 mM MgCl₂, 0.5% IGEPAL, 1 mM DTT, 0.5× protease inhibitor cocktail (Promega), 4 units/mL RNase inhibitor (Promega), and 100 µg/mL cycloheximide). Following a 10 min incubation on ice, embryos were mechanically lysed by passing the suspension six times through a 23G needle. The lysates were then clarified by 2 rounds of centrifugation at 21,000 × g for 10 min at 4°C. 500 μL of clarified supernatant was loaded onto a continuous 10–50% (w/v) sucrose gradient prepared in buffer containing 20 mM HEPES-KOH (pH 7.4), 100 mM KCl, 10 mM MgCl₂, 1 mM DTT, and 100 µg/mL cycloheximide. Gradients were centrifuged at 230,000 × g for 2.5 h at 4°C using an SW40 rotor. Polysome profiles were then analyzed by fractionating the gradients at 0.2 mL/min into 800 μL fractions using a BioComp gradient fractionator. Absorbance at 260 nm was monitored continuously during fractionation to assess RNA distribution.

For *ddx49* F0 knockout embryos, data were collected from 650 embryos (*n*=3). For WT samples, data were collected from 650 and 750 embryos, with experiments conducted across two independent days (*n*=5). The area under the curve (AUC) for peaks was calculated after baseline correction, with the baseline defined as the lowest absorbance value within fractions 2–12.

### Ddx49 cloning, expression and purification

cDNAs encoding wild-type zebrafish Ddx49 were PCR amplified and cloned into pJet, with site-directed mutagenesis used to generate a Ddx49-I224N corresponding to the *ssl* lesion, each confirmed by whole-plasmid sequencing. Ddx49 and Ddx49-I224N were then PCR-amplified from pJET-Ddx49 and pJET-Ddx49-I224N as templates and sub-cloned into pET28a and pEGFP-C1 vectors using BamHI and HindIII sites. The human GAPDH sequence was PCR-amplified using human cDNA as a template and cloned into the pEGFP-C1 vector using BamHI and XhoI sites. The details of the pGEX-DDX49 construct have been described (Awasthi et al., 2018). The expression and purification of GST-tagged human DDX49 protein were performed as described (Awasthi et al., 2018). The expression and purification of His-tagged Ddx49 and Ddx49-I224N proteins were performed as described (Verma et al., 2017).

### ATPase and helicase assays

ATPase and helicase activities of Ddx49, Ddx49-I224N and DDX49 were performed as detailed previously (Awasthi et al., 2018). 1 µM of each purified protein was used for ATPase assays, and the reaction mixtures were incubated at 37 °C for 2 hr. The amount of phosphates released in the ATPase assay was quantified by measuring the O.D. at 655 nm. For the helicase activity assay, 10-mer RNA labeled oligo (5′-GCU UUA CGG U-3′) labeled with Alexa Fluor 700 at 5′ end (Eurogentec) and the 40-mer RNA unlabeled oligo (5′-AAA AAC AAA ACA AAA CAA AAC AAA ACU AGC ACC GUA AAG C-3′) (Eurogentec) were annealed in the molar ratio of 1:1.5 and the annealed oligos were used as substrates. 0.5 µM of each protein was used for helicase assays, and the reaction mixtures were incubated at 37 °C for 1 hr. The final reaction mixtures were resolved on a 12% native polyacrylamide gel and imaged using the LI-COR Odyssey Dlx Imaging system.

### Circular dichroism (CD) spectroscopy

CD analyses were carried out in J-815 CD spectrometer (JASCO) using 3 µM of Ddx49, Ddx49-I224N and human DDX49 proteins as described previously (Verma et al., 2017).

### Sub-cellular localization and protein levels

Human 293T cells were procured from National Centre for Cell Science, India and cultured in DMEM supplemented with 10% FBS at 37 °C in 5% CO2. Transfections were performed by using polyethylenimine (Polysciences; PEI MAX; Cat No. 24765-1). For subcellular localization studies, 293T cells were seeded onto coverslips and transfected with GFP-Ddx49 or GFP-Ddx49-I224N constructs. After 36 hr, cells were fixed using 4% formaldehyde, stained with DAPI and embedded in mowiol mounting medium. Images of transfected cells were acquired using a 63X oil-immersion objective of a fluorescence microscope (Nikon Eclipse Ti). For protein level analyses, 293T cells were co-transfected with GFP-GAPDH and GFP-Ddx49 or GFP-Ddx49-I224N constructs. After 40 hr, cells were lysed, and lysates immunoblotted with GFP primary antibody (SCBT, Cat. No. sc-9996) and IRDye® 800CW Goat anti-Mouse IgG secondary antibody (Licorbio, Cat. No. 926-32210).

### Vasa staining

*ssl/ssl* and *ssl/+* siblings at 7-8 SSL were fixed overnight at 4°C in 4% paraformaldehyde. Specimens were incubated at room temperature for 1 hour in blocking solution composed of 5% NGS, 1x PBS, 0.2% Triton-X-100 and 0.01% sodium azide. We then incubated in 1:250 Genetex GTX128306 Vasa antibody (rabbit polyclonal) for two hours at room temperature. Specimens were incubated with and 1:200 anti-rabbit Alexa Fluor 568 (Life Technologies) overnight at 4°C, and were also DAPI stained. Images for counts were acquired on the Zeiss Axio Observer equipped with a Yokogawa CSU-X1M5000 laser spinning disc.

## Acknowledgements

Supported by NIH R35 GM122471 (D.M.P.), NSF MCB 2503218 (A.J.) and Department of Biotechnology, Government of India (Junior Research Fellowship (DBT-JRF) to A.K. Thanks to Katia Korzeniwsky for assistance with molecular cloning and all Parichy lab members for assistance with fish husbandry.

## Additional information

## Funding

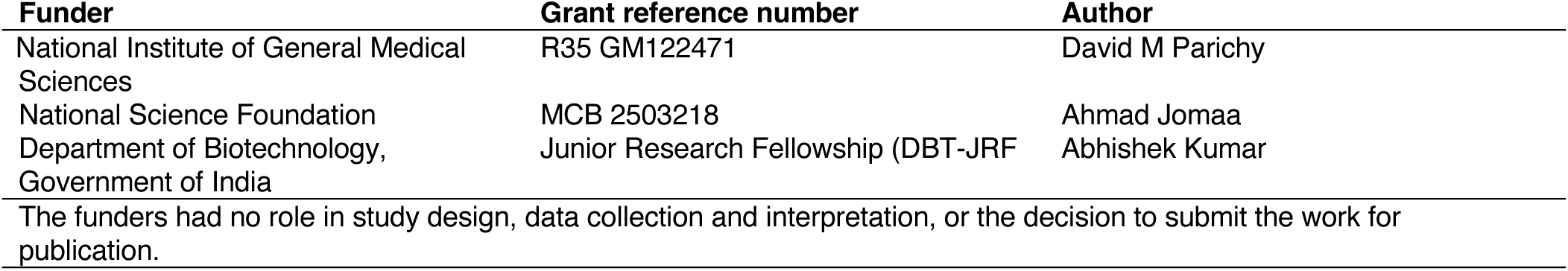

## Author contributions

Yipeng Liang, Conceptualization, Formal analysis, Investigation, Writing—review & editing; Kinga Malezyna, Investigation, Formal analysis; Abigail J Dennis, Investigation; Abhishek Kumar, Investigation; Braedan M McCluskey, Formal analysis; Diane S Sepich, Resources; Liliana Solnica-Krezel, Resources; Arunkumar Dhayala, Supervision; Ahmad Jomaa, Supervision, Funding acquisition; Larissa B Patterson, Conceptualization, Formal analysis, Investigation, Writing—review & editing; David M Parichy, Conceptualization, Writing—original draft, Writing—review & editing Visualization, Supervision, Project administration, Funding acquisition

## Ethics

This study was performed in strict accordance with the recommendations in the Guide for the Care and Use of Laboratory Animals of the National Institutes of Health. All of the animals were handled according to approved institutional animal care and use committee (IACUC) protocol (#4170) of the University of Virginia.

## Additional files

### Supplementary files

- Supplementary file 1. Supplementary tables
- Supplementary file 2. Figure source data

## Data availability

Sequencing data have been deposited in GEO under accession code GSE349762. The following dataset was generated.

| Author(s) | Year | Dataset title | Dataset URL | Database and Identifier |
| --- | --- | --- | --- | --- |
| Liang Y, Parichy D | 2026 | DEAD-box RNA helicase 49 (Ddx49) requirements for proliferative expansion of post-embryonic neural crest derived pigment cell populations and ribosome biogenesis | https://www.ncbi.nlm.nih.gov/geo/query/acc.cgi?acc=GSE349762 | NCBI Gene Expression Omnibus, GSE349762 |

